# SPIC-dependent erythrophagocytic macrophages drive granuloma formation and pathogen persistence during intracellular bacterial infection

**DOI:** 10.64898/2026.05.24.727563

**Authors:** Aaron Fountain, Weixian Lin, Max J. Lain, Yuan Xue, Trung H. M. Pham

## Abstract

Macrophages maintain tissue homeostasis by phagocytosing spent cells, recycling nutrients, and mounting antimicrobial responses to eliminate pathogens. Yet, they can also act as a cellular niche and organize granulomas that enable intracellular bacteria, such as *Salmonella enterica*, to persist in infected tissues. Here, using a murine *Salmonella* Typhimurium (*S*Tm) infection model, we find that granuloma formation and bacterial persistence are dependent on SPIC, which controls development of VCAM1^+^ macrophages critical for erythrocyte, heme, and iron recycling. VCAM1^+^ macrophages markedly increase in infected spleens and have high levels of erythrophagocytosis, intracellular bacteria, and T-cell co-stimulatory ligands. Using SPIC-deficient mice generated from CRISPR gene editing, we show that SPIC is required for macrophage co-stimulatory ligand expression and formation of a VCAM1^+^ macrophage zone that produces CXCL9 retaining T cells at the granuloma periphery. SPIC deletion abolishes this granuloma cellular architecture and reduces bacterial persistence. We propose that SPIC-dependent erythrophagocytic macrophages drive granuloma formation and bacterial tissue persistence.

## Introduction

Even with modern public health efforts, growing antibiotic arsenals, and vaccines, intracellular bacterial pathogens, including *Salmonella enterica*, *Mycobacterium tuberculosis* (*Mtb*), and *Bartonella henselae*, continue to burden hundreds of millions of people worldwide^1–3^. These pathogens can establish persistent infection and survive in infected tissues at low levels for long periods of time despite effector innate and adaptive antimicrobial immune responses and treatments^4–11^. Persistent infection presents substantial challenges due to perpetual pathogen reservoirs, lack of effective means to detect and monitor subclinical hosts, difficulty in eradicating low bacterial levels inside host cells, and prolonged antimicrobial therapies that drive antibiotic resistance^1,12^. Many antibiotics preferentially target replicating bacteria, thereby limiting their efficacy against persistent pathogens that are not actively growing^13,14^. Conventional vaccine designs elicit effector innate and adaptive immune mechanisms, yet these pathogens have evolved to persist even under such host immune pressure^8,15^. New insights into host and pathogen mechanisms underlying persistent infection may inform more effective therapeutic and vaccine strategies to combat these infections^16^.

Following infection at barrier sites, such as the gut, intracellular bacterial pathogens can be trafficked to lymphoid organs, including the draining lymph nodes, spleen, and bone marrow, which are enriched with mononuclear phagocytes^4,17–22^. The traffic of pathogens to lymphoid organs promotes host defense responses, such as myelopoiesis and induction of adaptive immunity. Intriguingly, intracellular pathogens may have evolved mechanisms to exploit this trafficking to establish persistence beyond barrier sites^23–28^. Although invasive tissue biopsy is not commonly performed in infected patients, available histopathological studies suggest lymphoid tissue persistence and granuloma formation are shared features in humans across infections with phylogenetically distinct intracellular bacteria^4,17,29,30^.

Granulomas are multicellular structures comprised of immune and stromal cells that form in tissues to contain infection^8,31–35^. Evidence suggests granuloma formation may also enable bacterial persistence^8,9,36^. Although considerable heterogeneity in immune cellular composition has been observed among individual granulomas in intracellular bacteria-infected mammalian tissues, lymphocytes typically concentrate around the granuloma periphery forming a lymphocytic cuff and are more sparse in the center^8,37^. In murine *Salmonella*-infected spleens, it was shown that despite robust Th1 cell activation and effector function required for bacterial restriction, CXCL9^+^/CXCL10^+^ mononuclear phagocytes form a zone surrounding the granuloma core that retains CXCR3^+^ Th1 cells and limits clearance of the bacteria that are localized in the granuloma center^8^. In *Mtb*-infected lungs of mice, humans, and non-human primates, IFNγ^+^ CD4 T-cells have also been found to be low in the center of granulomas, which correlate with diminished bacterial restriction^36–39^. Other findings suggest regulatory T cells may promote bacterial persistence in granuloma centers^38,39^. However, CXCR3 neutralization disrupting CXCL9^+^/CXCL10^+^ macrophage and Th1 cell interaction diminishes Th1 response and increases tissue bacterial burden^8^. Thus, we still lack a full understanding of how the cellular architectures of granuloma affect bacterial persistence and the underlying mechanisms.

Macrophages act as sentinels of tissue homeostasis and exhibit diverse functions^40^. Across steady state and inflamed tissues, macrophage populations are variably comprised of cells derived from embryonic precursors that seed tissues during development and from bone-marrow derived monocyte precursors throughout life^40^. Macrophages maintain homeostasis by phagocytosing spent cells, recycling nutrients, remodeling damaged tissues, and mediating effector innate immune responses to eliminate pathogens^40,41^. Yet, they can also act as a cellular niche and orchestrate granuloma formation that enables persistence of intracellular bacterial pathogens in infected tissues^5,8,9,41–45^. Recycling spent erythrocytes, heme, and iron is an essential tissue function of macrophages^46,47^. Classically, in mammalians, spent red blood cells are phagocytosed and removed from circulation by macrophages resident in the splenic red pulps and liver sinusoids, called red pulp macrophages and Kupffer cells, respectively^46,47^. These macrophages recycle iron required for most of the body’s daily iron needs. Increased presence of erythrophagocytic macrophages has been observed in different tissues under a variety of pathophysiological settings, including sepsis, intracellular bacterial infections, tissue damage, and the tumor microenvironment^47–51^. The biological relevance of erythrophagocytic macrophages may extend beyond iron homeostasis. In murine experimental models of lung and liver tissue damage, erythrophagocytic macrophages have been shown to dampen inflammation^50,52^. Intriguingly, during *Salmonella* infection, the pathogen may preferentially survive within macrophages that have engulfed erythrocytes^49,53^. A recent study suggests intracellular *Salmonella* is protected from iron deprivation by residing inside erythrophagocytic macrophages^49^.

The transcription factor (TF) SPIC is a member of the ETS TF family that includes SPI1 (PU.1) and SPIB^54,55^. Prior studies showed SPIC regulates development of erythrophagocytic macrophages in the spleen and liver^54,56^. Germline SPIC knockout mice have selective loss of red pulp macrophages and Kupffer cells expressing VCAM1, a transcriptional target of SPIC in macrophages and monocytes^50,52,54,56,57^. These knockout mice also have iron accumulation in the spleen and liver due to a defect in erythrocyte, heme, and iron recycling^54,56^. SPIC-expressing macrophages have since been observed in the bone marrow, thymus, tumor microenvironment, and in inflamed liver, gut, and lungs^50–52,57–59^. SPIC is induced in macrophages and monocytes by LPS, CpG, the alarmin IL-33, and heme, a tissue damage associated molecular pattern^52,57^. Very little is known about how SPIC and SPIC-dependent macrophages affect host immune responses during infections^60^.

Here, we seek to uncover mechanisms of macrophage functions underlying bacterial persistence in infected tissues using a persistent infection model with fully virulent *Salmonella* Typhimurium (*S*Tm) in 129x1/SvJ mice, which have intact NRAMP1 function required for efficient macrophage iron recycling and the host’s resistance to intracellular pathogens in humans and mice^61–64^. Comparative single-cell transcriptomics (scRNA-seq) analysis of macrophages and monocytes from uninfected and infected spleens at different stages of infection identifies a correlation between *Spic^+^Vcam1^+^* macrophage transcriptional state with establishment of persistent infection. Functional interrogation shows VCAM1^+^ macrophages in infected spleens have significantly higher levels of erythrophagocytosis, intracellular *S*Tm, and CD86, demonstrating that they are more primed to engage co-stimulatory receptors and activate T cells. SPIC deletion reduces VCAM1^+^, erythrophagocytic macrophage abundance, tissue bacterial persistence, and splenomegaly. We find in SPIC-deficient mice, macrophages in the spleens exhibit impaired CD86 expression and fail to form a distinct VCAM1^+^ macrophage zone at the granuloma periphery where CD4 T cells form a lymphocytic cuff. SPIC is required for CXCL9 expression in these macrophages and CD4 T cell retention at the lymphocytic cuff over time. Collectively, our findings suggest a model in which SPIC-dependent erythrophagocytic macrophages drive granulomatous response enabling persistent infection.

## Results

### Single-cell transcriptomics analysis identifies a correlation between SPIC-regulated macrophage transcriptional state with infection stage

To dissect mechanisms of macrophage functions and granuloma formation in intracellular bacteria-infected tissues, we use a persistent infection model with fully virulent *Salmonella* Typhimurium (*S*Tm) in 129x1/SvJ (SvJ) mice^5,6,9,41,42^. Susceptible mouse strains, such as C57BL/6, rapidly succumb to virulent *S*Tm^5^. In contrast, resistant strains, such as SvJ mice, can control the pathogen to low levels in infected tissues, including the spleen, and develop persistent infection lasting many months before clearing^5,8,65^. A genetic factor that has been linked to this susceptibility is *Slc11a1*^61,62^. Susceptible mice carry *Slc11a1^S^* variant encoding non-functional SLC11A1, a metal transporter critical for efficient iron and erythrocyte recycling in macrophages and host’s ability to control virulent intracellular pathogens^5,49,61–72^. Like most humans, SvJ and other mouse strains carry *Slc11a1^R^* and intact SLC11A1 function^5,65,68^. In this model, by 4 weeks post-inoculation (p.i.), we observe steady restriction of bacteria to low, persistent levels in infected spleens (Figure 1A), as persistent infection is being established^5,8,9^. Bonafide granulomas form that are histologically similar to granulomas observed in *Salmonella*-infected human tissues^4,5,9,73^.

**Figure 1:**
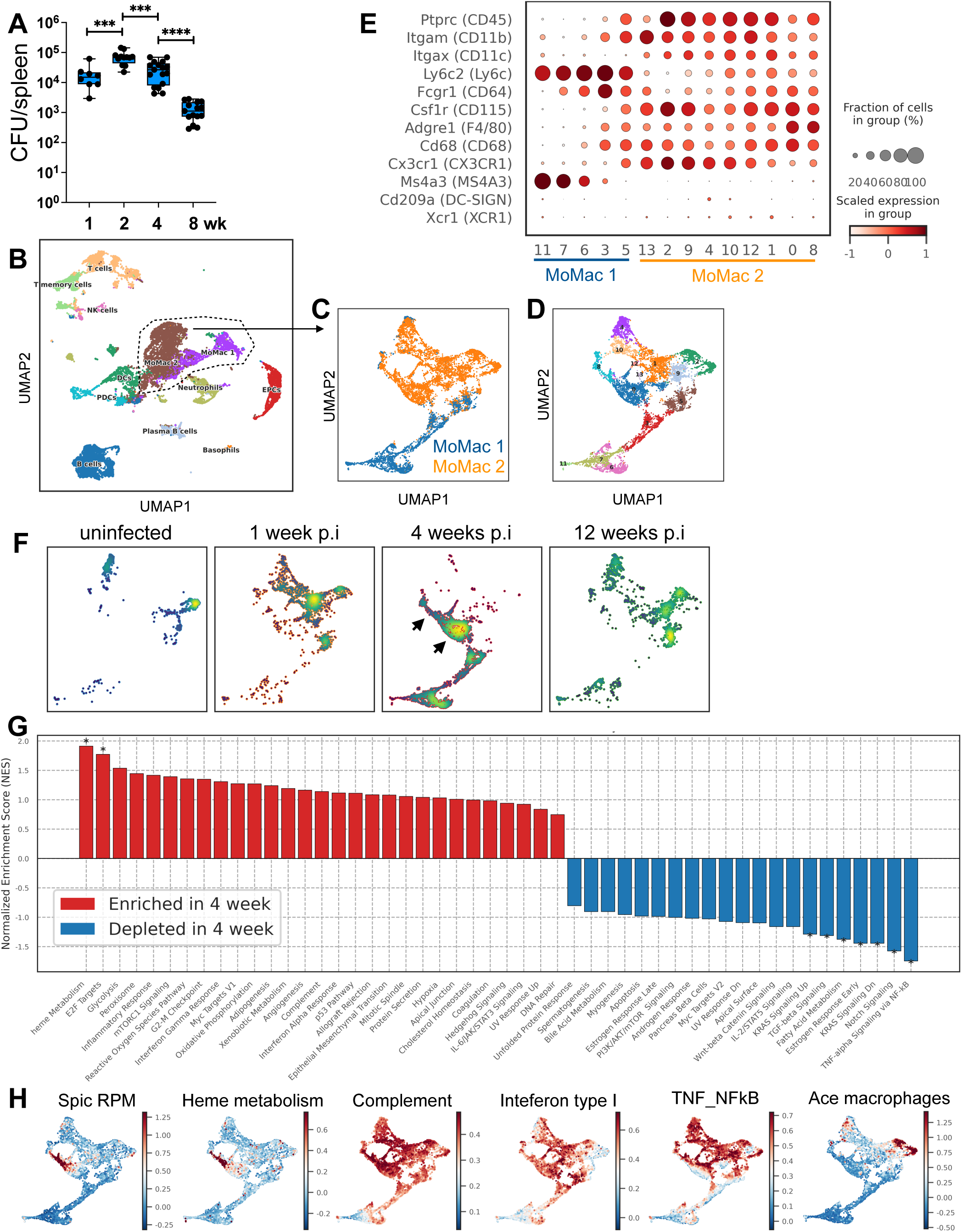
ScRNA-seq identifies a correlation between SPIC-regulated macrophages with infection stages. Mice infected with *S.* Typhimurium SL1344 via intraperitoneal inoculation and analyzed at indicated times p.i. (A) Quantitation of splenic bacterial burdens by colony forming units (CFU) using plating assays. Bars: geometric mean. Dots: individual mice. Significance calculated using a two-tailed Mann-Whitney test. n ≥ 8 mice per time point. (B) UMAP projection of cells from scRNA-seq of FACS-enriched splenocytes from uninfected and infected spleens at 1, 4, and 12 weeks p.i. (see Methods). Label abbreviations: monocytes and macrophages (MoMac), dendritic cells (DCs), plasmacytoid dendritic cells (pDCs), natural killer cells (NK cells), erythroid precursor cells (EPCs). (C) UMAP projection of the MoMac 1 and MoMac 2 clusters (D) UMAP projection of the distinct transcriptional states among MoMac 1 and MoMac 2 cells. (E) Dotplot showing expression levels of myeloid marker genes among MoMac 1 and MoMac 2 transcriptional states. (F) Relative abundances of distinct transcriptional states projected in (D) at different stages of infection. Arrowheads indicate two clusters, 0 and 8 in (D), that are enriched in 4 weeks p.i. (G) Pathway, gene set enrichment (GSEA) analysis comparing sequenced cells from 4 weeks p.i. and cells combined from uninfected, 1 week, and 12 weeks p.i. (H) Pathway Activity score of immune response gene sets using top differentially expressed genes (see Methods).

We applied 10X Genomics single-cell RNA-sequencing (scRNA-seq) to identify macrophage functional states and pathways influencing bacterial tissue persistence. In prior studies, we had characterized granuloma-associated CD11b^+^CD11c^+^Ly6C^+^ macrophages (hereafter macrophages) in infected spleen that are distinct from classical monocytes (CM), classical dendritic cells (cDC), and CD11b^+^CD11c^+^ monocytes (Figure S1)^9,42^. These cells express phenotypic and functional markers consistent with activated macrophages^9,42^. Furthermore, they express inducible nitric oxide synthase (iNOS) and harbor intracellular *S*Tm at a markedly higher frequency than other myeloid cells in infected spleens^9,42^. We devised a FACS-enrichment strategy that enriches for a full spectrum of these macrophages, their precursors, and simultaneously captures other immune cells in infected spleens for scRNA-seq (Figure S1 and Methods)^42^. We sequenced samples from uninfected spleens and infected spleens at 1, 4, and 12 weeks p.i. to delineate critical cell states and functional pathways involved in different infection stages, from acute to long-term persistent infection.

As expected, our scRNA-seq dataset is enriched in macrophages and monocytes (MoMac) and includes other immune cell populations in infected spleens (Figure 1B). Within MoMac, a subset of MoMac 1 population (clusters 11, 7, and 6) have high expression of *Ms4a3*, a marker of the granulocyte-monocyte progenitor, and low expression of differentiated monocyte and macrophage markers, suggesting progenitor cell states (Figure 1C-E)^42,74^. The MoMac 2 clusters (0, 1, 2, 4, 8, 9, 10, 12, 13) exhibit phenotypic and functional markers consistent with macrophages and monocytes (Figure 1E). Distinct transcriptional states are enriched at different stages of infection (Figure 1F). Cluster 8 and 0, for instance, are more enriched at 4 weeks p.i., suggesting they may play a role in establishing the persistent infection stage. To gain insights into the biological pathways differentially enriched at this infection time point, we performed gene set enrichment analysis (GSEA) using the publicly available Molecular Signature Database (MSigDB) gene modules^75^. For this analysis, we excluded the progenitor states with high *Ms4a3* levels (MoMac 1 clusters 11, 7, and 6), which have high numbers of cell-cycle related genes^42^. We grouped the remaining cluster transcriptomes from 4 weeks p.i. into one group and compared them to those from uninfected, 1 week, and 12 weeks p.i. combined (Figure 1F). This analysis revealed that heme metabolism pathways are relatively enriched in macrophages and monocytes at 4 weeks p.i. (Figure 1G), suggesting that heme metabolism function in these cells may play a role in establishing persistent infection stage.

Increasing evidence suggests macrophages that uptake, metabolize, and recycle erythrocytes, heme, and iron can modulate inflammatory response in damaged tissues^47,50,76^. Intriguingly, erythrophagocytic macrophages have been observed to act as a cellular niche for *S*Tm and enable bacterial survival^49,53,77^. Thus, erythrophagocytic and heme metabolizing macrophages may impact host immune responses and infection persistence. We next sought to identify the cell populations among the MoMac 2 clusters (Figure 1C-D) that are differentially enriched in heme metabolism genes by performing Pathway Activity Score Analysis (Materials and Methods). This analysis reveals cells with the highest composite expression of genes in the heme metabolism gene module. We also included a SPIC Macrophages gene module (*Spic*, *Vcam1*, *Hmox1*, *MerTK*, *Hba-a1*, and *Alas2*) that we and others have used to define VCAM1^+^ erythrophagocytic macrophages in the spleen and other tissues, and complement pathway module, which we previously observed to have similar enrichment pattern as SPIC macrophage module^42,50,51^. In addition, we incorporated gene modules for Interferon Type I Response, TNF_NFKb Signaling, and ACE Macrophages that we observed in prior studies to be more enriched in cell states that are distinct from SPIC-expressing macrophages^42^. Our Pathway Activity Score Analysis showed that the SPIC macrophage and heme metabolism gene modules are both highly enriched in cluster 0 and 8, which are more abundant at the 4-week infection stage (Figure 1D, F, H). The Complement gene module is relatively enriched in these cell clusters, but to a lesser extent. In contrast, expression of the ACE Macrophages module is enriched in cluster 2 and 4 (Figure 1H). These findings suggest cluster 0 and 8 represent SPIC-regulated macrophages involved in erythrocyte and heme recycling in infected tissues and their functional states correlate with a stage when persistent infection is being established.

### VCAM1^+^ macrophages exhibit high erythrophagocytic and co-stimulatory capacity

Very little is known about the role of SPIC in macrophage responses during infection^60^. SPIC has been reported to be expressed in macrophages, monocytes, and B cells^54,55^. In our scRNA-seq datasets from *S*Tm-infected spleens, *Spic* is highly expressed in macrophages and monocytes (MoMac 2 clusters), and to a much lesser extent in neutrophils and dendritic cells (Figure 2A). *Vcam1* is a transcriptional target of SPIC^54,56,57^. *Spic* and *Vcam1* expressions are highly enriched and concordant in MoMac cluster 0 and 8 (Figure 2B). These cells also express high levels of functional markers of erythrophagocytic, heme recycling macrophages, including *Hmox1*, *Slc40a1*, *MerTK,* and *C1q* (Figure 2B)^50,56,78^. To functionally interrogate *Spic^+^Vcam1^+^* macrophages in infected tissues, we stained VCAM1 on myeloid cells among splenocytes from *S*Tm-infected spleens at 4 weeks p.i. We prepared splenocyte single-cell suspension and subjected them to erythrocyte lysis before staining (see Methods). We find that VCAM1 expression is highest on macrophages among myeloid cells in infected spleens (Figure 2C, S1).

**Figure 2:**
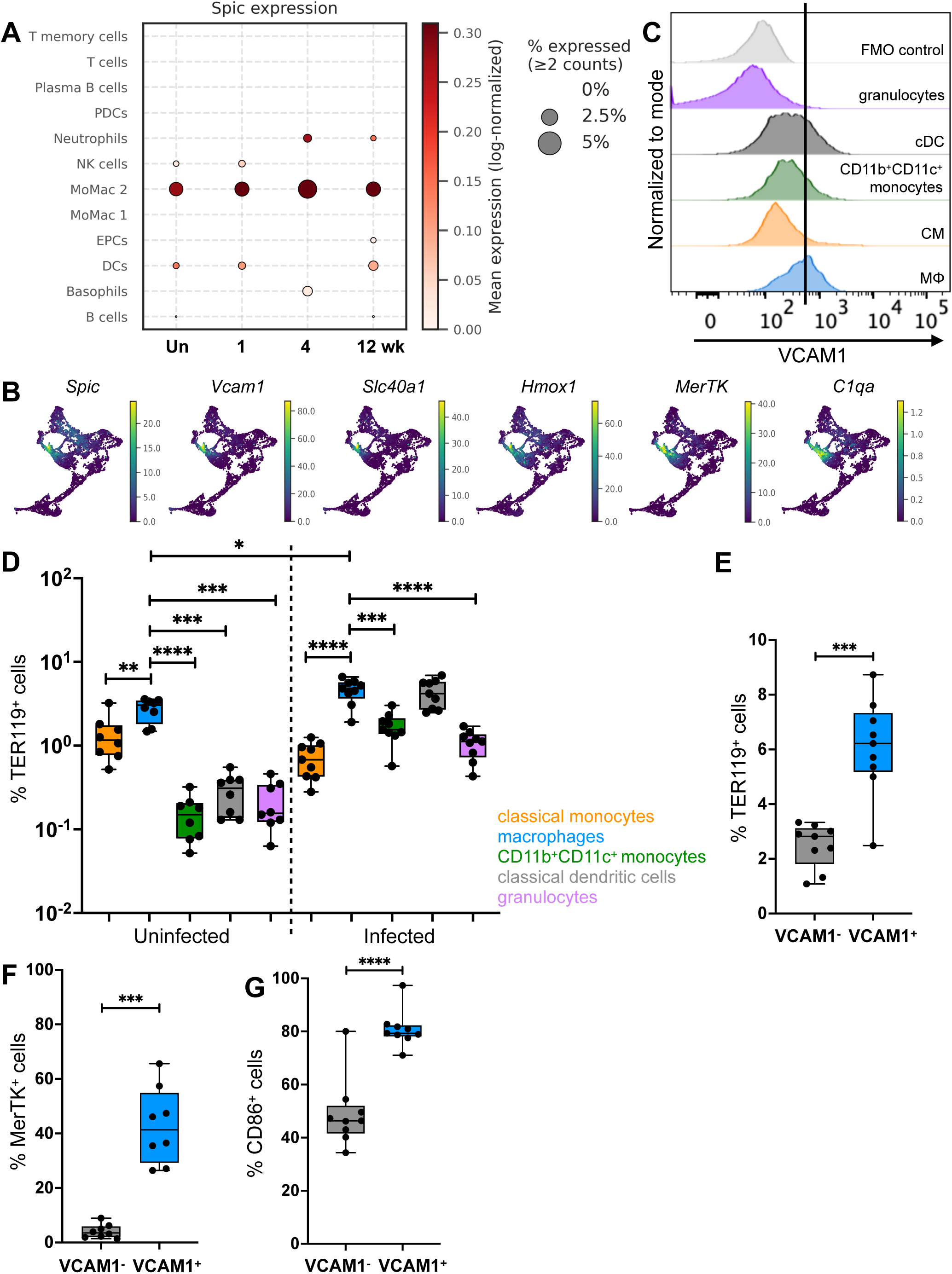
VCAM1^+^ macrophages exhibit high erythrophagocytic and co-stimulatory capacity. (A) Mean *Spic* expression among different cell types captured in our scRNA-seq. (B) Expression of phenotypic markers associated with *Spic^+^Vcam1^+^*erythrophagocytic macrophages among the MoMac cell clusters from scRNA-seq shown in Figure 1D. (C) Flow cytometric analysis of splenic macrophages and myeloid cells as gated in Figure S1. Histograms illustrating VCAM1 expression on macrophages (MΦ), classical monocytes (CM), CD11b^+^CD11c^+^ monocytes, classical dendritic cells (cDC), and granulocytes. (D) Frequencies of TER119^+^ cells quantitated using intracellular staining and flow cytometric analysis among splenic myeloid cells. Myeloid cell populations are gated as shown in Figure S1. Dots: individual mice. (E) VCAM1^-^ and VCAM1^+^ macrophages are gated as in Figure S1. Frequencies of intracellular TER119^+^ cells among VCAM1^-^ and VCAM1^+^ macrophages by flow cytometry shown. (F) Frequencies of MERTK^+^ cells among VCAM1^-^ and VCAM1^+^ macrophages by flow cytometry. Gating example shown in Figure S2A. (G) Frequencies of CD86^+^ cells among VCAM1^-^ and VCAM1^+^ macrophages by flow cytometry. Gating example shown in Figure S2B. Significance calculated using a two-tailed Mann-Whitney test. C-G, n ≥ 9 mice, from at least 2 independent experiments.

We next assessed the erythrophagocytic capacity of VCAM1^+^ macrophages in infected spleens by staining these cells and other myeloid cells for intracellular TER119^+^ erythrocytes using intracellular staining^79,80^. In both uninfected and infected spleens, macrophages stained positive for intracellular TER119 at higher frequencies, compared to CM, granulocytes, and CD11b^+^CD11c^+^ monocytes, with a median frequency of 5% in infected spleens (Figure 2D and S2A). Classical dendritic cells have markedly lower TER119^+^ frequencies, compared to macrophages, in uninfected spleens, but their frequencies rise to similar level as macrophages in infected spleens (Figure 2D). Importantly, the TER119^+^ frequencies of VCAM1^+^ macrophages are more than 2-fold higher, compared to VCAM1^-^ macrophages (Figure 2E). Consistent with our transcriptional analysis of *Spic^+^Vcam1^+^* macrophages (Figure 2B), VCAM1^+^ macrophages also have high levels of MerTK, a TAM family tyrosine kinase and efferocytotic receptor expressed abundantly on erythrophagocytic macrophages (Figure 2F, S2A)^42,51^.

Prior studies showed that splenic dendritic cells and macrophages that have engulfed erythrocytes upregulate activation marker and T cell co-stimulatory ligand CD86 and exhibit a more activated profile^81–83^. To test if VCAM1^+^ macrophages in *S*Tm-infected spleens, which have high frequencies of erythrophagocytosis are more activated, we measured CD86 levels. We found that VCAM1^+^ macrophages express significantly higher CD86, compared to VCAM1^-^macrophages (Figure 2G, S2C). Collectively, these findings indicate that VCAM1^+^ macrophages have distinctly high level of erythrophagocytosis and capacity to stimulate adaptive cellular immunity in infected spleens.

### VCAM1^+^ erythrophagocytic macrophages markedly increase in infected spleen and harbor bacteria during persistent infection

To gain further insights into the role of VCAM1^+^ macrophages in infected spleen, we examined the population dynamics of these cells over the course of infection. We quantitated VCAM1^+^ macrophage numbers by flow cytometry and found that this cell population increased by ∼200-fold by 4 weeks p.i. (Figure 3A). Corresponding to the marked increase of VCAM1^+^ macrophages, the numbers of TER119^+^ macrophages increased by 260-fold (Figure 3B), while the increase in TER119^+^ CM was more modest at 40-fold (Figure 3C). We performed Perl’s Prussian blue stain to functionally correlate erythrocyte recycling in the infected spleens with the drastic increase in erythrophagocytic macrophages and myeloid cells. This histochemical stain detects ferric iron and highlights hemosiderin, a breakdown product of erythrocytes, in the splenic red pulp^54^. In uninfected spleens, the stain signal was robustly detected in the red pulp (Figure 3D). In contrast, the stain signal was significantly diminished in infected spleens at 4 weeks p.i. These findings suggest increased erythrophagocytosis activity during persistent infection.

**Figure 3:**
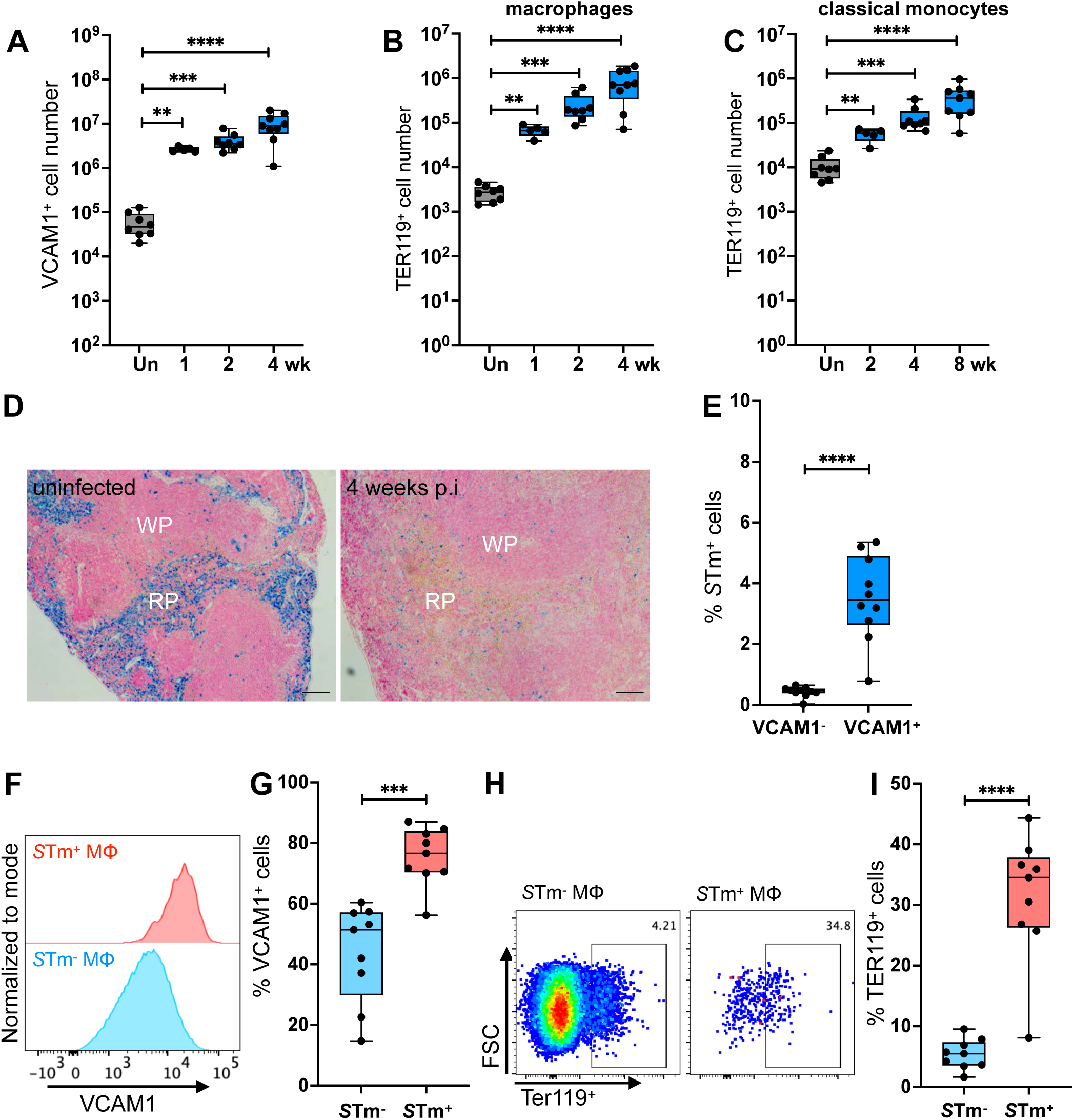
VCAM1^+^ erythrophagocytic macrophages markedly increase and harbor bacteria during persistent infection. (A-C) Splenocytes from uninfected and infected mice analyzed by flow cytometry at indicated time points p.i. Gating of myeloid cell populations is shown in Figure S1. (A) Numbers of splenic VCAM1^+^ macrophages. (B) Numbers of splenic TER119^+^ macrophages. C) Numbers of splenic TER119^+^ classical monocytes. Dots: individual mice. (D) Perl’s Prussian Blue stain for ferric iron on formalin-fixed spleens from uninfected and infected mice at 4 weeks p.i. WP: white pulp. RP: red pulp. Scale bar: 200 μm. (E-I) Infected mice analyzed at 4 weeks p.i. (E) Frequencies of intracellular *S*Tm^+^ cells among VCAM1^-^ and VCAM1^+^ macrophages from infected spleens at 4 weeks p.i. quantitated by flow cytometry. *S*Tm levels measured by fluorescent signal of Tomato-expressing *S*Tm or by intracellular staining using fluorescent anti-*Salmonella* antibody. Dots: individual mice. (F) Splenic macrophages analyzed by flow cytometry were gated for *S*Tm^+^ and *S*Tm^-^. Histogram show VCAM1 expression on *S*Tm^+^ vs. *S*Tm^-^ macrophages. (G) Frequencies of VCAM1^+^ cells among *S*Tm^+^ and *S*Tm^-^macrophages. Dots: individual mice. (H) Splenic macrophages analyzed by flow cytometry were gated for *S*Tm^+^ and *S*Tm^-^, then visualized for intracellular TER119 signal as shown. (I) Frequencies of TER119^+^ cells among *S*Tm^+^ and *S*Tm^-^ macrophages. Dots: individual mice. Significance calculated using a two-tailed Mann-Whitney test. A-C, E, G, I, n ≥ 8 mice per group from at least 2 independent experiments.

*S*Tm has been observed to survive within macrophages that have engulfed erythrocytes^49,53,77^. However, our finding that VCAM1^+^ erythrophagocytic macrophages express more CD86 activation marker suggests these macrophages may be more activated to mount antimicrobial responses. Thus, we assessed whether VCAM1^+^ erythrophagocytic macrophages harbor intracellular bacteria to enable persistent infection. We infected mice with either Tomato^+^ *S*Tm and measured Tomato signal, or wildtype *S*Tm and stained for intracellular bacteria using anti-*Salmonella* antibody^8,9,24,42^. We observed that VCAM1^+^ macrophages have significantly higher *S*Tm^+^ frequencies, compared to VCAM1^-^ macrophages (Figure 3E). Furthermore, when we compared *S*Tm^+^ macrophages and *S*Tm^-^ macrophages in the infected spleens, we observed that *S*Tm^+^ macrophages have markedly higher VCAM1 levels and TER119^+^ frequencies (Figure 3F-I). Collectively, our results demonstrate that infection induces VCAM1^+^ erythrophagocytic macrophage abundance in the spleen. These macrophages are primed to activate adaptive cellular immunity but harbor bacteria that may contribute to persistent infection.

### SPIC deletion accelerates pathogen clearance and lessens splenomegaly

To determine the impact of SPIC on host immune response and infection, we leveraged CRISPR gene editing technologies to create germline deletion of SPIC in SvJ mice. We designed gRNAs to target the coding regions of exon 4 and exon 6 and successfully induced a 7-nucleotide insertion and 2-nucleotide substitution causing a premature STOP codon at amino acid 48 of SPIC (Figure S3A) (see Methods). We confirmed the loss-of-function allele (hereafter, knockout (KO) allele) by sequencing. For routine genotyping, we developed a qRT-PCR assay using hydrolysis probes that differentially hybridize and discriminates the *Spic* wildtype (WT) and KO alleles (Figure S3B)^84^. Western blotting showed loss of SPIC expression (Figure S3C). Similar to *Spic^-/-^* C57BL/6 mice generated from homologous recombination gene targeting reported previously, the new SPIC KO SvJ mice we have generated are born at lower than expected Mendelian ratio, but are fertile and appear healthy^54^.

We infected SPIC WT and KO mice with *S*Tm and analyzed at 4 weeks p.i. to determine the role of SPIC in macrophage responses and host immunity. We found that the VCAM1 median fluorescent intensity (MFI) and the VCAM1^+^ frequencies of macrophages in infected spleens were significantly reduced in KO mice (Figure 4A-B). VCAM1 expression on CM was low and there was a statistically significant reduction in KO CM, compared to WT CM (Figure 4A-B). We found that the total numbers of VCAM1^+^ macrophages decreased by more than 5-fold in KO mice (Figure 4C) by 4 weeks p.i. These findings are consistent with the role of SPIC in the development of VCAM1^+^ macrophages^54,56,57^. Persistent *S*Tm infection causes drastic splenomegaly in humans and mice^4,5,85^. Remarkably, the spleen sizes of infected SPIC KO mice were significantly smaller, compared to WT mice (Figure 4D). By weight, KO spleens were ∼4-fold less than WT spleens (Figure 4D-E). Concurrently, we observed a 0.5 log reduction in bacterial burdens in the spleens of KO mice (Figure 4F).

**Figure 4:**
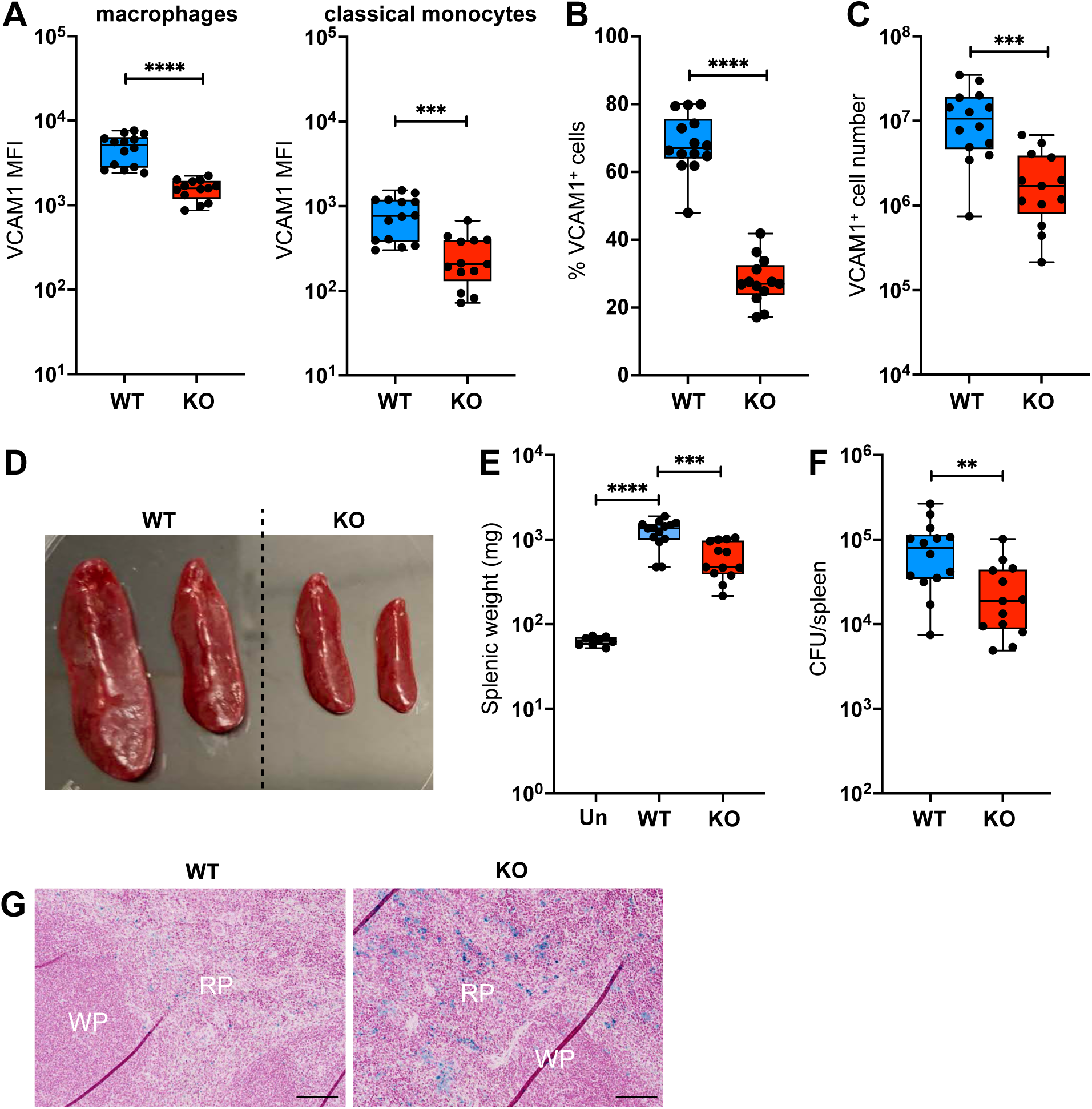
SPIC deletion accelerates pathogen clearance and lessens splenomegaly. Infected mice analyzed at 4 weeks p.i. Splenocytes analyzed by flow cytometry and gated for macrophages and other myeloid cells as shown in Figure S1. (A) Median fluorescent intensity (MFI) of VCAM1 on total macrophage and classical monocyte populations. (B) Frequencies of VCAM1^+^ macrophages in SPIC WT and KO spleens. (C) Total VCAM1^+^ macrophage cell numbers in SPIC WT and KO spleens. (D) Example of reduced splenomegaly in SPIC KO mice. (E) Splenic weights of uninfected (Un) and infected SPIC WT and KO mice. (F) Bacterial levels in infected SPIC WT and KO spleens quantitated by CFU plating assay. (G) Perl’s Prussian Blue stain for ferric iron on formalin-fixed spleens from infected SPIC WT and KO mice. WP: white pulp. RP: red pulp. Scale bar: 100 μm. Significance calculated using a two-tailed Mann-Whitney test. A-G, data from 3 independent experiments 4-5 WT and KO mice per experiment.

We performed Perl’s Prussian blue staining for erythrocyte-associated ferric iron in the splenic red pulp to assess how SPIC deletion affects erythrocyte recycling and iron distribution in infected spleens. We found that KO mice show stronger Prussian Blue staining in the splenic red pulp, compared to WT mice. Increased erythrophagocytic macrophages have been observed in inflamed and *S*Tm-infected liver^49,50^. To test if SPIC has a similar functional impact in infected liver as the spleen, we also measured bacterial burdens and performed Prussian blue stain on liver samples. We observed a significant reduction in bacterial levels and stronger ferric iron staining in SPIC KO compared to SPIC WT infected livers (Figure S4A and S4B). Collectively, these results demonstrate that SPIC is required for VCAM1^+^ erythrophagocytic macrophage function in infected tissues during persistent infection, which hampers pathogen clearance.

### SPIC is required for macrophage expression of co-stimulatory ligands and bacterial persistence over time

We next investigated the mechanism underlying accelerated bacterial clearance resulting from SPIC deletion. Previous studies reported that SPIC function dampens inflammatory cytokine production, including TNF and IL-6, in macrophages and SPIC/VCAM1^+^ erythrophagocytic macrophages act as anti-inflammatory macrophages in tissue damage settings^50,52,59^. Thus, we tested if SPIC exerts similar effects on macrophage inflammatory response in infected tissues. We obtained total splenocytes from infected SPIC WT and KO mice at 4 weeks p.i., stimulated them *ex-vivo* with heat-killed *S*Tm (HKST) as PAMP stimuli, and measured intracellular cytokine levels. We found that SPIC WT and KO macrophages from infected spleens have similar TNF and IL-6 levels (Figure S5A-B), suggesting that the accelerated bacterial clearance we observed in SPIC KO mice was not driven by heightened macrophage TNF and IL-6 response. Furthermore, we found that the numbers of granulocytes, CM, activated IL-18R^+^ CD4 T cells, and IL-18R^+^IFNγ^+^ CD4 T cells were all lower in infected spleens of KO mice, compared to WT mice (Figure S5C-I). Collectively, these findings suggest that heighted inflammatory cellular responses were not the cause of the accelerated bacterial clearance in SPIC KO animals.

We then examined whether SPIC deletion reduces the abundance of erythrophagocytic macrophage niche that harbors bacteria enabling persistent infection. We found that the frequencies of intracellular TER119^+^ macrophages were only slightly lowered in SPIC KO spleens at 4 weeks p.i. (Figure S5J). However, the numbers of TER119^+^ macrophages decreased by 5-fold in the SPIC KO spleens compared to WT spleens, indicating a significant reduction of the erythrophagocytic macrophage niche (Figure 5A). Among the remaining macrophages, the frequencies of *S*Tm^+^ macrophages were similar in the infected spleens of SPIC WT and KO mice (Figure 5B).

**Figure 5:**
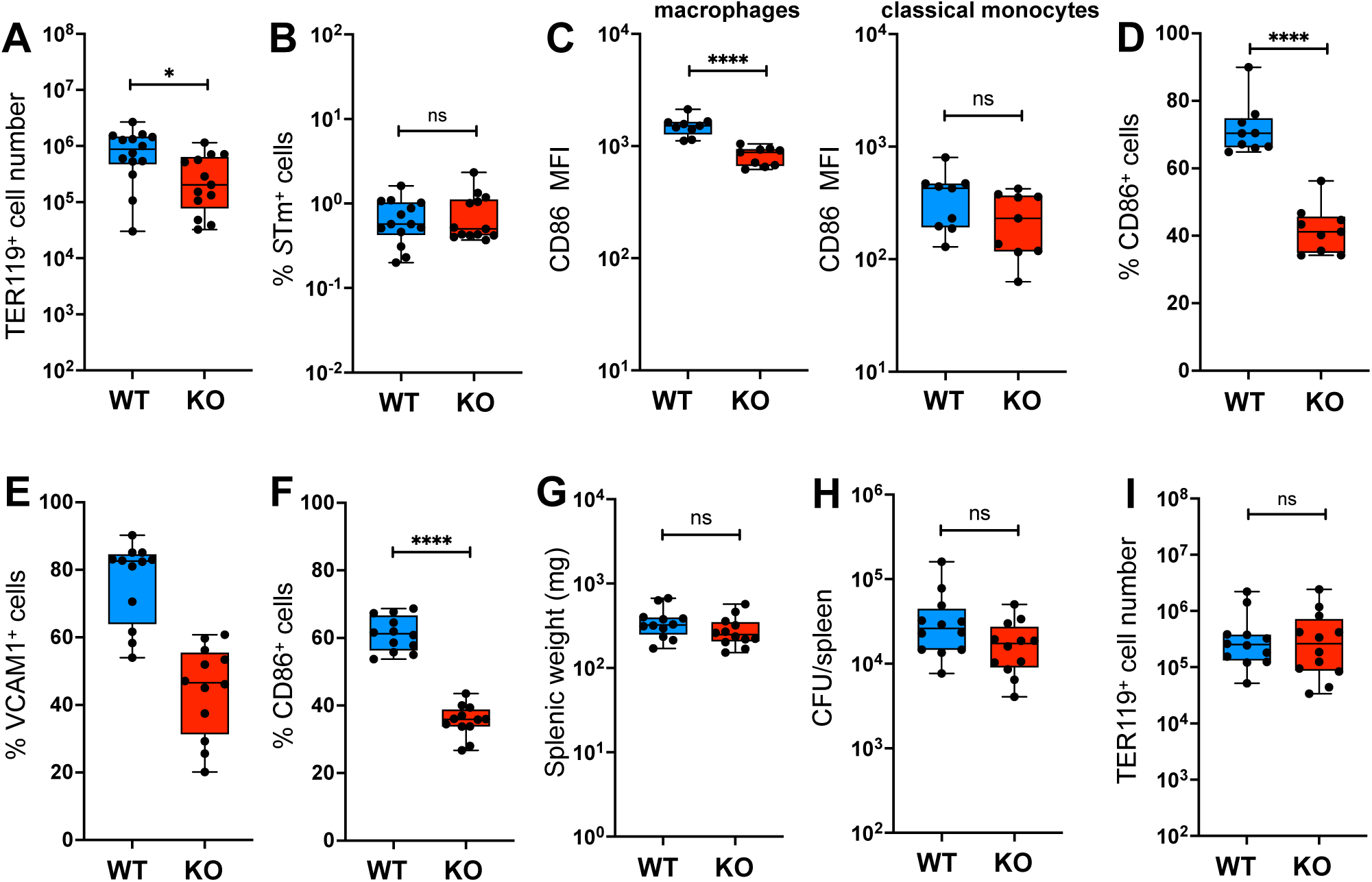
SPIC is required for macrophage co-stimulatory ligand expression and bacterial persistence over time. (A-D) Infected mice analyzed at 4 weeks p.i. Splenocytes analyzed by flow cytometry and gated for macrophages and other myeloid cells as shown in Figure S1. (A) Numbers of TER119^+^ macrophages in SPIC WT and KO spleens. (B) Frequencies of *S*Tm^+^ macrophages in WT and KO spleens. (C) CD86 median fluorescent intensity (MFI) of macrophages and classical monocytes in SPIC WT and KO spleens. (D) Frequencies of CD86^+^ macrophages in infected SPIC WT and KO mice. (E-I) Infected mice analyzed at 2 weeks p.i. Splenocytes analyzed by flow cytometry and gated for macrophages and other myeloid cells as shown in Figure S1. (E) Frequencies of VCAM1^+^ macrophages in the infected spleens of SPIC WT and KO mice. (F) Frequencies of CD86^+^ macrophages. (G) Splenic weights of SPIC WT and KO mice at 2 weeks p.i. (H) Splenic CFU in SPIC WT and KO spleens at 2 weeks p.i. (I) Frequencies of TER119^+^ macrophages in infected WT and KO spleens. Significance calculated using a two-tailed Mann-Whitney test. A-I, data from 3 independent experiments 4-5 WT and KO mice per experiment.

Our observations that VCAM1^+^ erythrophagocytic macrophages have higher CD86 and intracellular bacterial levels demonstrate an association between erythrophagocytosis and bacterial exposure with macrophage activation and co-stimulatory capacity. To assess whether SPIC and macrophage CD86 expression are linked, we measured CD86 levels on myeloid cells from infected spleens of SPIC WT and KO mice at 4 weeks p.i. Strikingly, we found that CD86 levels, as measured by median fluorescence intensity (MFI), were markedly reduced in SPIC KO macrophages (Figure 5C). In contrast, CD86 expression was low in CM and there was no significant difference between SPIC WT and KO CM (Figure 5C). When compared by frequencies of CD86^+^ cells, the median frequency of CD86^+^ SPIC WT macrophages was 70% compared to 41% for KO macrophages (Figure 5D and S2C). Collectively, our observations suggest that the impaired bacterial persistence in the infected spleens of KO mice is not due to heightened inflammatory cellular immune responses. Rather, reduced bacterial persistence may lead to lower macrophage activation and effector cellular immune responses.

To gain further insights into the mechanisms of SPIC, we infected mice and performed analysis earlier at 2 weeks p.i., when bacterial levels are yet to be under control in infected spleens (Figure 1A), to delineate the sequence of SPIC-dependent effects that hamper bacterial clearance in persistent infection. Like our observations at 4 weeks p.i., we found that there was a marked reduction of VCAM1^+^ and CD86^+^ macrophages in *S*Tm-infected spleens of SPIC KO mice, compared to WT mice at this infection stage (Figure 5E-F). However, there was no significant differences in splenomegaly, tissue bacterial burden, or TER119^+^ macrophage abundance at this time point (Figure 5G-I). These findings suggest SPIC is required for activation and co-stimulatory capacity of macrophages, development of VCAM1^+^ macrophages, and intriguingly, infection persistence over time.

### SPIC deficiency impairs macrophage CXCL9 production and formation of granuloma macrophage-CD4 T cell interaction zone

The dependency of macrophage CD86 expression on SPIC prompted us to consider whether SPIC affects macrophage-T cell interaction and the *in vivo* context in which this pathway functions. In our prior study, we observed that VCAM1^+^ macrophages were more detectable at the granuloma periphery and diminished in the center^42^. Others have found that Th1 cells typically concentrate at the periphery of granulomas in part due to CXCL9 and CXCL10 chemoattractant signal produced by a group of macrophages surrounding granulomas^8^. This CXCL9/CXCL10-mediated retention has been proposed to limit Th1 cell infiltration and promote eradication of bacteria localized at the granuloma centers. These observations led us to hypothesize the SPIC-dependent VCAM1^+^ macrophages, which have robust CD86 levels, produce CXCL9 and/or CXCL10 required for macrophage-Th1 cells interaction at the granuloma periphery. To interrogate this hypothesis, we leveraged our scRNA-seq data to assess CXCL9 and CXCL10 expression in *Spic^+^Vcam1^+^* macrophages. We discovered that these macrophages robustly express CXCL9 and CXCL10 (Figure 6A).

**Figure 6:**
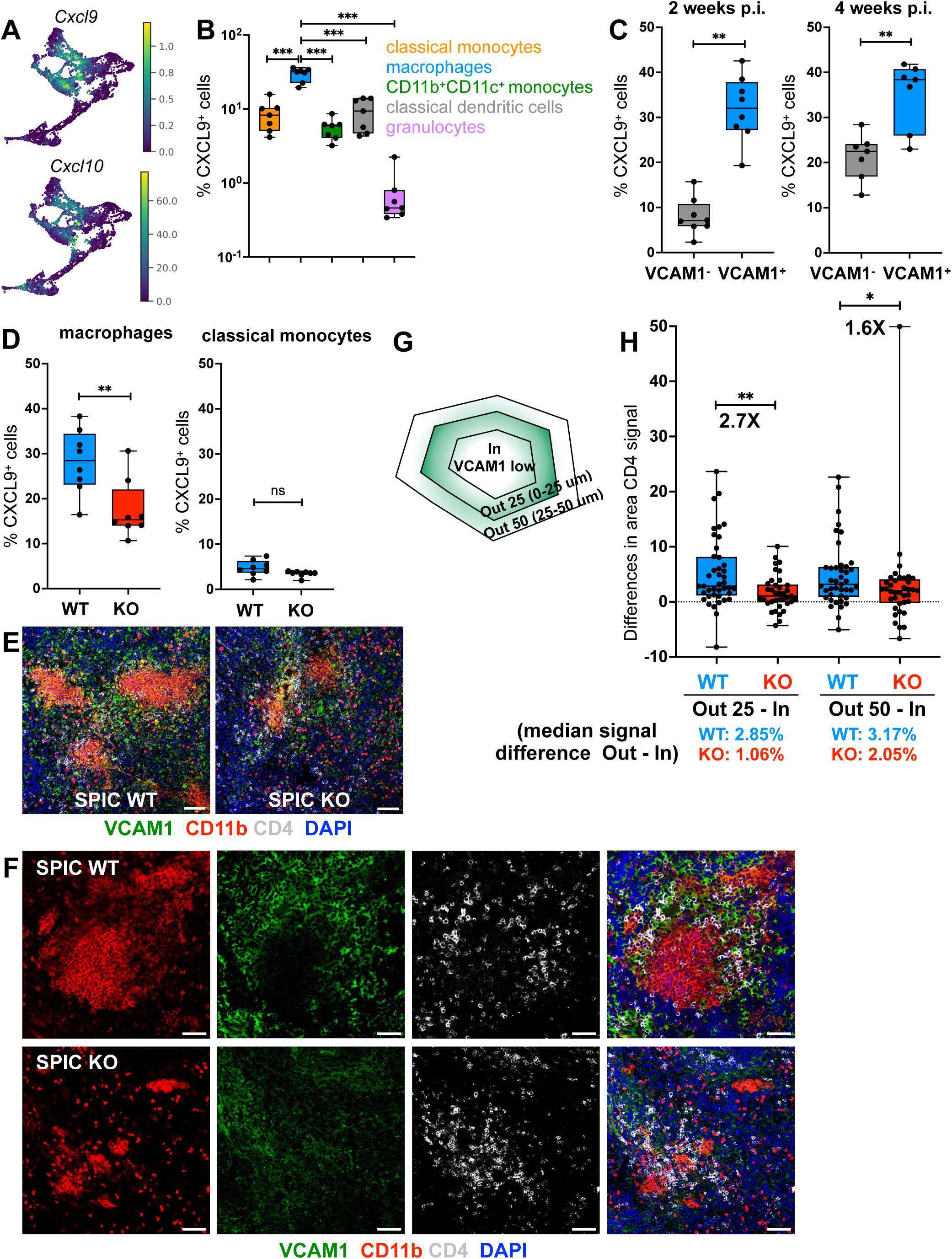
SPIC deficiency impairs macrophage CXCL9 production and formation of granuloma macrophage-CD4 T cell interaction zone. (A) Expression of *Cxcl9* and *Cxcl10* among the MoMac cell clusters from scRNA-seq as in Figure 1D. (B) Frequencies of CXCL9^+^ cells among different myeloid populations infected SPIC WT spleens at 4 weeks p.i. quantitated by flow cytometry analysis. Dots: individual mice. (C) Frequencies of CXCL9^+^ cells among VCAM1^-^ and VCAM1^+^ macrophages in infected SPIC WT spleens at 2 and 4 weeks p.i. (D) Frequencies of CXCL9^+^ macrophages and classical monocytes (CM) in infected SPIC WT and KO spleens at 2 weeks p.i. quantitated. (E) Staining of VCAM1^+^CD11b^+^ and CD4 T cells in nascent granulomas at 2 weeks p.i. (F). Co-localization of VCAM1^+^CD11b^+^ and CD4 T cells in granuloma areas in SPIC WT and KO spleens at 4 weeks p.i. (G) A diagram of regions in the granuloma center and outer zones used for quantitation of CD4 T cells signal. (H) Median differences in the CD4 T cell staining signal between areas depicted in (G). Dots: individual quantified granulomas. Significance calculated using a two-tailed Mann-Whitney test. A-C, n ≥ 8 mice per group. E-H, n ≥ 4 mice.

We then used flow cytometry to quantitate CXCL9 expression in VCAM1^+^ macrophages and other myeloid cells in infected spleens. We found that a median of 32% of VCAM1^+^ macrophages are positive for CXCL9, at least 3-fold higher than the median frequencies in other myeloid cells (Figure 6B). Furthermore, CXCL9^+^ frequencies among VCAM1^+^ macrophages are significantly higher than VCAM1^-^ macrophages (Figure 6C). Since VCAM1^+^ macrophages are dependent on SPIC, we tested if SPIC deletion affects the levels of CXCL9-producing macrophages. We found that by 2 weeks p.i., the frequencies of CXCL9^+^ macrophages were significantly lowered in the infected spleens of SPIC KO mice, compared to WT animals. In contrast, CXCL9 expression was low in CM and there was no significant difference between SPIC WT and KO CM (Figure 6D). These findings indicate that VCAM1^+^ macrophages are a key SPIC-dependent CXCL9-producing cell population in infected spleens.

Next, we investigated how the loss of VCAM1^+^ and CXCL9^+^ macrophages in the spleens of SPIC KO mice affects granuloma macrophage-T cell interaction and formation using confocal microscopy. We previously found that the pan-myeloid cell marker CD11b stains *S*Tm granulomas robustly and we use this marker to demarcate granuloma areas from the surrounding^9,33,42^. By 2 weeks p.i., we observed that the presence of VCAM1^+^ macrophages in the outer zone of nascent granulomas is reduced and many granulomas are mildly diminished in KO spleens (Figure 6E). By 4 weeks p.i. in SPIC WT spleens, many granulomas have a distinct VCAM1^+^ macrophage-CD4 T cell zone at the granuloma periphery, forming a lymphocytic cuff surrounding the granuloma core. In contrast, in KO spleens granulomas are often diminished and lack a distinct VCAM1^+^ macrophage-T cell zone, with more granulomas showing increased T cell infiltration into the granuloma center (Figure 6F and S6). We quantitated the loss of CD4 T cell cuff across granuloma images from a group of WT and KO mice. To do this, we used open-source FIJI analysis software. We masked CD4 signal of the images and used a combination of CD11b and VCAM1 signal to draw a boundary around the granuloma core where VCAM1 signal is lower (**In**side area) (Figure 6G). A 25-μm concentric area was drawn around the **In**side boundary, 0 to 25-μm away, and designated as **Out 25** area (Figure 6G). A second concentric area, 25 to 50-μm from the **In**side boundary, was also drawn and designated as **Out 50** area (Figure 6G). The area covered by the CD4 signal as a proportion of the **In**side, **Out 25**, and **Out 50** areas were individually calculated to create the proportions that quantify CD4 signal coverage. We found that the median difference of CD4 signal between **Out 25** area to the **In**side was 2.85% in the SPIC WT spleens (Figure 6H). In the KO spleens, the median difference between the **Out 25** area to **In**side area or granulomas was 1.06%. When compared, there is a 2.7-fold reduction in median difference between granulomas in the KO versus WT spleens. When the median difference of the CD4 signal was calculated between the **Out 50** area and **In**side area, the median difference was only 1.6-fold less for the SPIC KO, compared to WT granulomas (Figure 6H). Thus, the SPIC-dependent positioning of CD4 T cells is strongest in the area immediately adjacent to the granuloma center. Taken together, our results suggest that SPIC is required for formation of a VCAM1^+^ macrophage - CD4 T cell interaction zone at the granuloma periphery and loss of this granuloma cellular architecture is associated with impaired bacterial persistence.

## Discussion

By leveraging a persistent *Salmonella* infection model, newly created SPIC SvJ knockout mice, and a multimodal approach, we defined SPIC as a determinant of granuloma formation and persistent intracellular bacterial infection. Building on prior research, we found that SPIC promotes development of VCAM1^+^, erythrophagocytic macrophages in *S*Tm-infected spleens. These macrophages have high levels of intracellular TER119^+^ erythrocytes, *S*Tm, and T-cell co-stimulatory ligand CD86, suggesting that they are activated to stimulate T cell responses. Remarkably, SPIC germline deletion reduces bacterial burdens in the spleens and livers, instead of enabling bacterial growth. SPIC KO mice have similar tissue bacterial levels at 2 weeks p.i. but lower bacteria levels at 4 weeks compared to WT mice, indicating that SPIC function promotes persistent infection. Mechanistically, SPIC deletion reduces the abundance of VCAM1^+^, erythrophagocytic macrophages in the spleen that can harbor bacteria during persistent infection. Furthermore, SPIC is required for macrophage CD86 and CXCL9 expression driving formation of a VCAM1^+^ macrophage-T cell interaction zone that retains T cells at the granuloma periphery (Figure 7). Our findings suggest a model in which SPIC-dependent erythrophagocytic macrophages orchestrate cellular architecture that drives granuloma formation and persistent intracellular bacterial infection.

**Figure 7.**
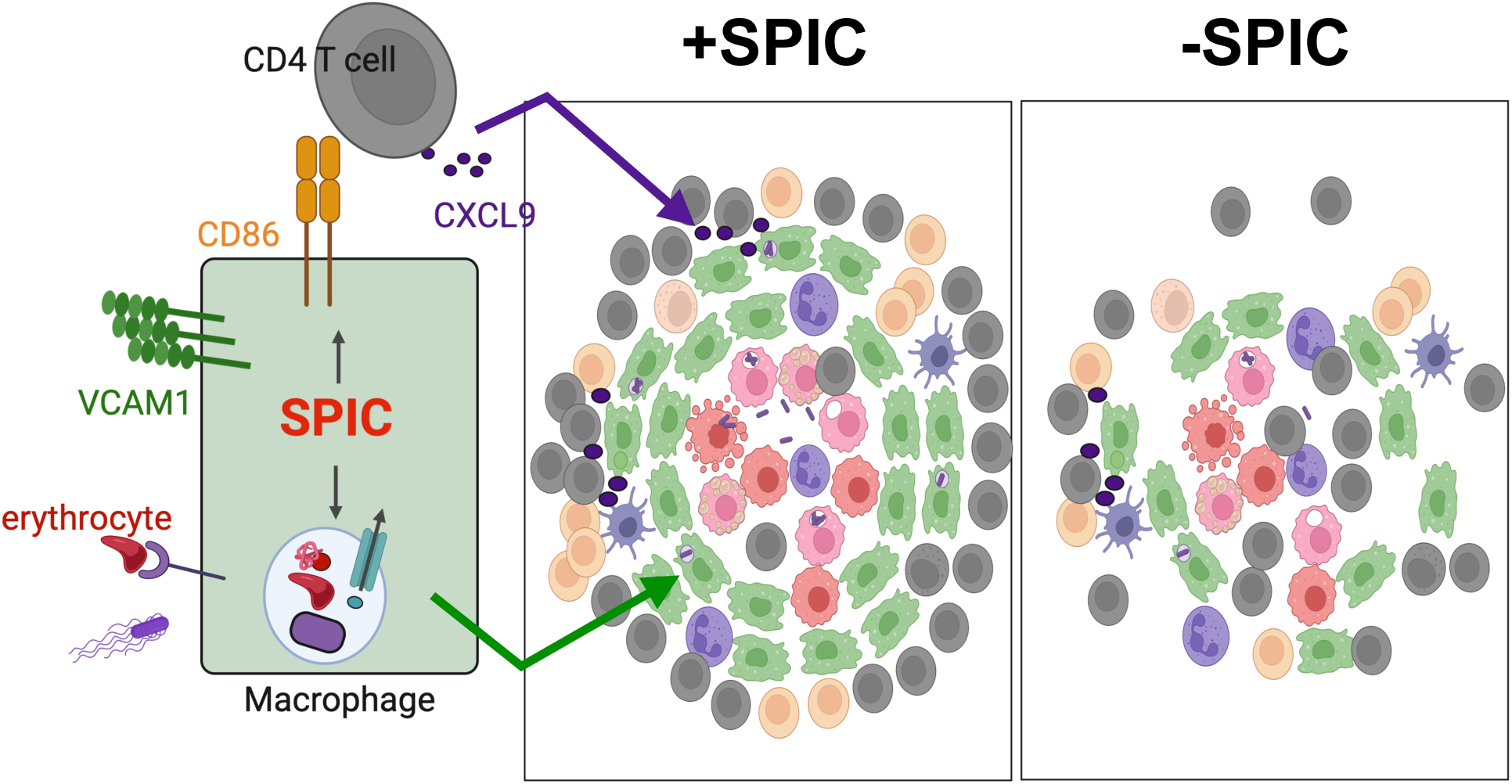
Proposed model for SPIC-dependent erythrophagocytic macrophages driving granuloma formation and bacterial persistence in *Salmonella*-infected tissues: Infection and spent erythrocytes in infected tissues trigger induction of SPIC, which is required for expression of macrophage CD86 and development of VCAM1^+^ erythrophagocytic macrophages. These macrophages localize to the outer zone of granulomas, provide co-stimulatory signals to T cells via CD86, and produce CXCL9 that retains CXCR3^+^ CD4^+^ T cells at the granuloma periphery forming a lymphocytic cuff. SPIC deletion impairs formation of this VCAM1^+^ macrophage-T cell interaction zone and granuloma cellular architecture accelerating bacterial clearance.

Findings from HIV-infected patients and T-cell depleting studies in animals infected with *M. tuberculosis* and *S. enterica* demonstrate that CD4 T cell immunity is required for bacterial restriction in these infections^8,86–88^. A conundrum, then, is how the pathogens can persist in infected tissues for periods of time despite robust effector CD4 T cell response. It has been hypothesized that the concentration of CD4 cells around the granuloma periphery, known as a lymphocytic cuff, reduces the presence of these cells and limits eradication of bacteria in the granuloma core^8,37^. However, direct evidence supporting this hypothesis has been lacking. In persistent *S.* Typhimurium infection in mice, the retention of CXCR3^+^ CD4 T cells in the lymphocytic cuff is mediated by CXCL9/CXCL10-producing macrophages localized in the granuloma periphery^8^. Disruption of this macrophage-CD4 T cell interaction using CXCR3 neutralizing antibody increased bacterial burden, instead of enhancing bacterial clearance. This perturbation also interferes with their effector function critical for bacterial restriction, making it challenging to assess how formation of the lymphocytic cuff and granuloma cellular architecture contribute to bacterial persistence. Our findings in this study provide evidence that formation of the VCAM1^+^ macrophage-T cell interaction zone and lymphocytic cuffs at the granuloma periphery promotes bacterial persistence during *S*Tm infection. Mechanistically, this granuloma cellular architecture is dependent on SPIC. Phylogenetically distinct intracellular bacteria, such as *S. enterica* and mycobacteria, have different life cycles and microbial pathogenesis strategies. However, host granulomatous response to contain these pathogens in infected tissues may share common underlying mechanisms. It is possible that the positioning of CXCR3^+^ CD4 T cells in the lymphocytic cuff observed across different host species and intracellular bacterial granulomas commonly requires the presence of CXCL9/CXCL10-producing macrophages analogous to the SPIC-dependent macrophages we observe here^37,89^.

Our results also suggest that the VCAM1^+^, erythrophagocytic macrophages play a unique role in shaping infection outcome by providing persistent T-cell stimulation within the granuloma microenvironment that drive granuloma formation and bacterial persistence over time. Consistent with this notion, we observed defects in macrophage CD86 and CXCL9 expression at 2 weeks p.i. when nascent granulomas develop, but no significant differences in bacterial levels or splenomegaly emerged yet. By 4 weeks p.i., when granulomas become cohesive aggregates of macrophages and other immune cells with a more distinct VCAM1^+^ macrophage zone and T cell cuffs, differences in bacterial levels and splenomegaly are apparent. Persistent stimuli are thought to be a key element of granulomatous response^34^. The proximity of VCAM1^+^ macrophages with T cells at the lymphocytic cuff, their infected state, and high CD86 and CXCL9 expression suggest they may function as local antigen presenting cells to drive continued T cell response and granuloma formation, restraining the pathogen within the granuloma microenvironment during persistent infection. In line with this hypothesis, treatment with anti-CD4 or CXCR3 neutralizing antibodies in an infected host during persistent, controlled infection disrupted bacterial control and increased bacterial burdens^8,88^. On the other hand, VCAM1^+^, erythrophagocytic macrophages hamper bacterial elimination by retaining CD4 T cells at the granuloma periphery and harboring persisting bacteria. Disruption of the VCAM1^+^ macrophage-T cell interaction zone at the granuloma periphery in SPIC-deficient animals may expose more bacteria-harboring cells, including the iNOS^+^ macrophages in the granuloma cores, to T cell-mediated killing^8^. SPIC deletion, overall, led to enhanced bacterial clearance, indicating that effector antimicrobial innate and adaptive immune mechanisms remain sufficient to provide immunity against the pathogen in the absence of SPIC function. Whether intracellular pathogens, such as *S. enterica*, have evolved pathogenesis mechanisms to strategically exploit this unique macrophage population remain to be explored. Conceptually, our findings suggest modulating host pathways, such as SPIC, and their regulatory networks represents a unique strategy to combat pathogens capable of establishing persistent infection despite robust effector innate and adaptive immune response.

SPIC-expressing, erythrophagocytic macrophages have been observed in a variety of steady state and inflamed tissues, including spleen, liver, thymus, bone marrow, gut, lung, and tumor microenvironment^50–52,54,58,59,90^. These macrophages carry out critical biological functions including phagocytosing spent erythrocytes and heme-containing proteins to recycle iron, maintaining oxygenation capacity essential for metazoan life, and minimizing tissue-damaging heme excess^47,50,54,57^. Our understanding of erythrocyte, heme, and iron recycling is evolving. Emerging evidence suggests hemolysis occurs as part of the erythrocyte recycling process not only in special scenarios, such as in the setting of hemolytic anemia, but also during steady-state erythrocyte turnover and inflammation^76^. Heme, which is a product of erythrocyte breakdown and a DAMP, the alarmin IL-33, as well as PAMPs such as LPS and CpG, can potently induce SPIC in monocytes and macrophages^52,56,57^. These upstream signals of the SPIC pathway may present together in an infected and damaged tissue. Future studies dissecting how these signals individually and interactively affect SPIC functions, macrophage tissue damage response, effector functions, and immunoregulation of T cell responses in immune microenvironments such as granulomas and tumors will provide important insights to advance our understanding infection and immunity.

## Acknowledgements

The authors thank for Drs. Susan Brewer and Denise Monack for critical reading of the manuscript. We are grateful to Drs. Denise Monack at Stanford University and Marc Jenkins at the University of Minnesota for sharing bacterial strains. Research reported in this publication was supported by grant K08-AI143796 from the NIAID (THMP), the Stanford Department of Pediatrics, the Maternal and Child Health Research Institute (THMP), the Pediatric Infectious Diseases Society Fellowship Award (THMP), and the Pediatric Infectious Diseases Society SUMMERS Award (MJL).

The content is solely the responsibility of the authors and does not necessarily represent the official views of the National Institutes of Health.

## Author contributions

Conceptualization A.F. and T.H.M.P.; Investigation: A.F., W.L., M.J.L., Y.X., and T.H.M.P.; Software management: YX; Project management: A.F. and T.H.M.P.; Research supervision: T.H.M.P.; Writing — Original Draft, A.F. and T.H.M.P.; Writing — Review & Editing, all authors; Funding Acquisition: T.H.M.P.

## Materials and Methods

### Lead Contact and Materials Availability

Further information and requests for resources and reagents should be directed to and will be fulfilled by the Lead Contact, Trung H. M. Pham

### Experimental Model and Subject Details

#### Ethics Statement

Experiments involving animals were performed in accordance with NIH guidelines, the Animal Welfare Act, and US federal law. All animal experiments were approved by the Stanford University Administrative Panel on Laboratory Animal Care (APLAC) and overseen by the Institutional Animal Care and Use Committee (IACUC). Animals were housed in a centralized research animal facility accredited by the Association of Assessment and Accreditation of Laboratory Animal Care (AAALAC) International.

#### Mouse Strains and Husbandry

129X1/SvJ mice were obtained from Jackson Laboratories or in-house SPIC KO 129X1/SvJ colonies. Male and female mice (7-16 weeks old) were housed under specific pathogen-free conditions in filter-top cages that were changed bi-monthly by veterinary or research personnel. Sterile water and food were provided *ad libitum*. Mice were given at least one week to acclimate to the Stanford Animal Biohazard Research Facility prior to experimentation.

#### Bacterial Strains and Growth Conditions

*Salmonella enterica* serovar Typhimurium strain SL1344 (gift from Dr. Denise Monack, Stanford University) and SL1344 *Tomato* (gift from Dr. Marc Jenkins, University of Minnesota)^8^ were used for this study. For all mouse infections, *S.* Typhimurium strains were maintained aerobically on Luria-Bertani (LB) agar supplemented with 200 μg/mL streptomycin and grown aerobically to stationary phase overnight at 37 °C with broth aeration. Bacterial cultures were spun down and washed with sterile phosphate-buffered saline (PBS) before suspension in PBS for infection.

### Methods Details

#### Generation of SPIC knockout mice using CRISPR-Cas9 gene-editing

*Spic* knockout (KO) gRNAs were designed to target the coding regions of exon 4 and exon 6. Ribonucleoprotein (RNP) complex of gRNA and Cas9 protein (IDT, Montana, USA) was prepared at final concentrations of 32 ng/μL gRNA and 50 ng/μL Cas9 protein. Female 129x1/SvJ mice purchased from Jackson Laboratory were superovulated with 5.5 units of PMSG followed by 5.5 units of hCG and were then mated with 129x1/SvJ males. Zygotes were collected and microinjected with the RNP complex into the pronucleus, followed by implantation into the oviducts of CD1 surrogate mothers. The resulting pups were genotyped to identify *Spic*-KO founder mice. A founder with 7–nucleotide insertion and 2–nucleotide substitution causing a premature stop codon at amino acid 48 was selected for further validation. To verify SPIC KO at the protein level, mouse spleen, liver, and lung were homogenized in RIPA containing HALT protease inhibitor and PMSF using a TissueRuptor II homogenizer. Lysates were analyzed with Western Blot using SPIC (PA106516, Thermofisher). This mouse line was confirmed to be a SPIC knockout mouse line.

Mice were genotyped using the qPCR method as described previously^84^. Briefly, DNA within crude ear tissue lysates was amplified in PCR using an outer pair of primers flanking the indel region. The initial PCR product was then diluted 1:1000 and analyzed using multiplex qPCR amplification on QuantStudio 3 (Applied Biosystems), whereupon an inner pair of primers amplified the diluted PCR product, and hydrolysis probes were used to report the presence of WT or SPIC indel sequences.

Primers and probes were synthesized by IDT (San Jose, CA), with hydrolysis probes synthesized as a PrimeTime Probe. For the outer primer pairs, the forward primer sequence is 5’-CCGTAGAGGGAATGGGTTATG-3’ and the reverse primer sequence is 5’-CTTCCTCTCTCTGCTGCTATTT-3’. For the inner primer pairs, the forward primer sequence is 5’-GGTTGCTTACAGATGTGTTCATA-3’ and the reverse primer sequence is 5’-TGACATTCCATAGTAGTTAGCG-3’. The PrimeTime probe for WT is 5’-TATCCTCACGTCAGAGGCAA-3’ modified with FAM and Zen/Iowa Black Fluorescent Quencher; the PrimeTime probe for the Indel is 5’-CAATCCGTACAGAGGCAACG-3’ modified with SUN and Zen/Iowa Black Fluorescent Quencher. Data was analyzed Design and Analysis 2 (DA2) software (Applied Biosystems).

#### Mouse Infections

For experiments involving only WT mice, age and sex matched mice were allocated to control and experimental groups randomly, sample sizes were chosen based on previous experience to obtain reproducible results and the investigators were not blinded. For experiments involving SPIC KO and WT mice were matched by age and sex to the KO mice wherever possible. Mice were inoculated intraperitoneally (*i.p.*) with 1-2 x 10^3^ CFU *S.* Typhimurium SL1344 or SL1344 *tomato* in 200 μL PBS. Mice were euthanized at the indicated time points post-inoculation by CO2 asphyxiation followed by cervical dislocation as the secondary method of euthanasia. Organs were collected, weighted, and either homogenized in PBS for colony-forming unit (CFU) enumeration, used to make single cell suspension for flow cytometric analysis, or prepared for microscopy examinations.

#### Flow Cytometry

Spleens from mice were minced with surgical blades No. 22 and incubated in digestion buffer (HBSS+Ca^2+^+Mg^2+^ + 50 μg/mL DNase (Roche) + 25 μg/mL Liberase TL (Sigma)) at 37 °C for 25 min, mixing at 200 rpm. EDTA was added at a final concentration of 5 mM to halt digestion. Single cell suspensions were passed through a 70-μm filter and washed with R5 buffer (RPMI containing 5% FBS and 10 mM Hepes). Red blood cells were lysed with ACK Lysis Buffer (Fisher Scientific) for 3 min at room temperature, washed, and resuspended in R5 buffer until they were stained for flow cytometry.

For myeloid intracellular cytokine staining, heat-killed S. Typhimurium (HKST) was prepared from stationary-phase broth cultures grown aerobically overnight followed by heat treatment for 30 min at 56 °C. Digested splenocytes were washed and resuspended in RPMI containing 10% FBS, 10mM Hepes, and 50 ug/mL gentamicin. A total of 20 x 10^6^ splenocytes in 2-mL volume was added into each well of 6-well plates and allowed to equilibrate for 30 min at 37 °C. HKST at MOI 10:1 and brefeldin A to final concentration of 3 μg/mL were then added to stimulate splenocytes. After 3 hours of stimulation at 37 °C, splenocytes were harvested and stained.

Single-cell suspensions were incubated in Fc Block (TruStain fcX anti-mouse CD16/32, Biolegend) for 15 min on ice and washed with PBS. Cells were stained on ice for 30 min in PBS with a cocktail of Live/Dead Fixable Blue Viability Dye (Invitrogen) and antibodies for surface antigens (list of antibodies used provided in the Key Resources table 1). Cells were washed with FACS buffer (PBS containing 2% FBS and 2 mM EDTA), followed by fixation for 15 min with Cytofix/Cytoperm solution (BD Biosciences). Cells were washed twice with Perm/Wash buffer (BD Biosciences) and stained for intracellular *Salmonella*, TNF, iNOS, CXCL9, IL-6, TER119, and IFNγ. For experiments with Typhimurium SL1344 *tomato,* splenocytes from mice infected with Typhimurium SL1344 were included for staining control. After washing, cells were resuspended in FACS buffer and analyzed on FACSymphony A5 analyzers (Becton Dickinson). Data were acquired with DIVA software (BD Biosciences) and analyzed using FlowJo software (BD Biosciences).

**Table 1.**
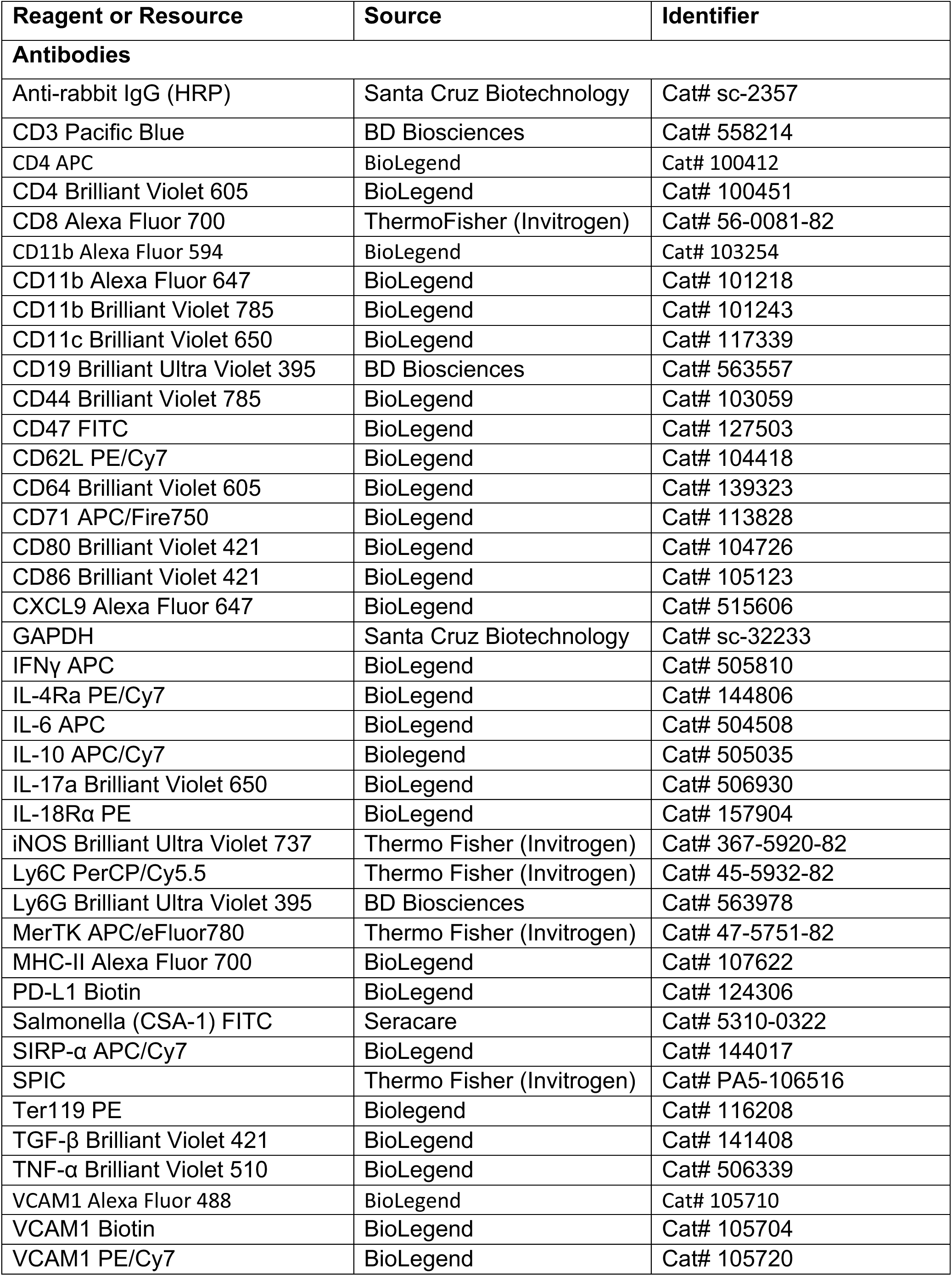
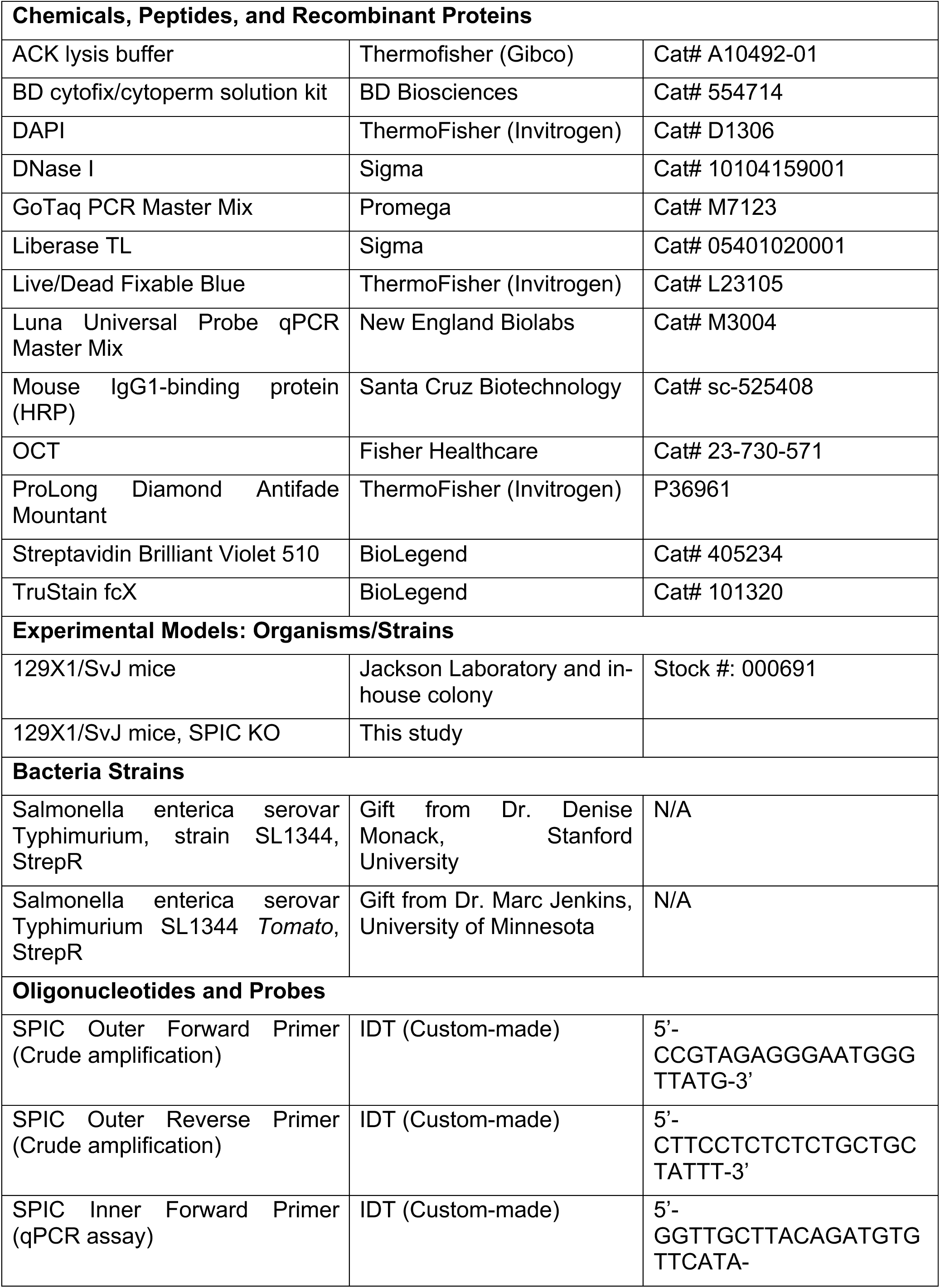
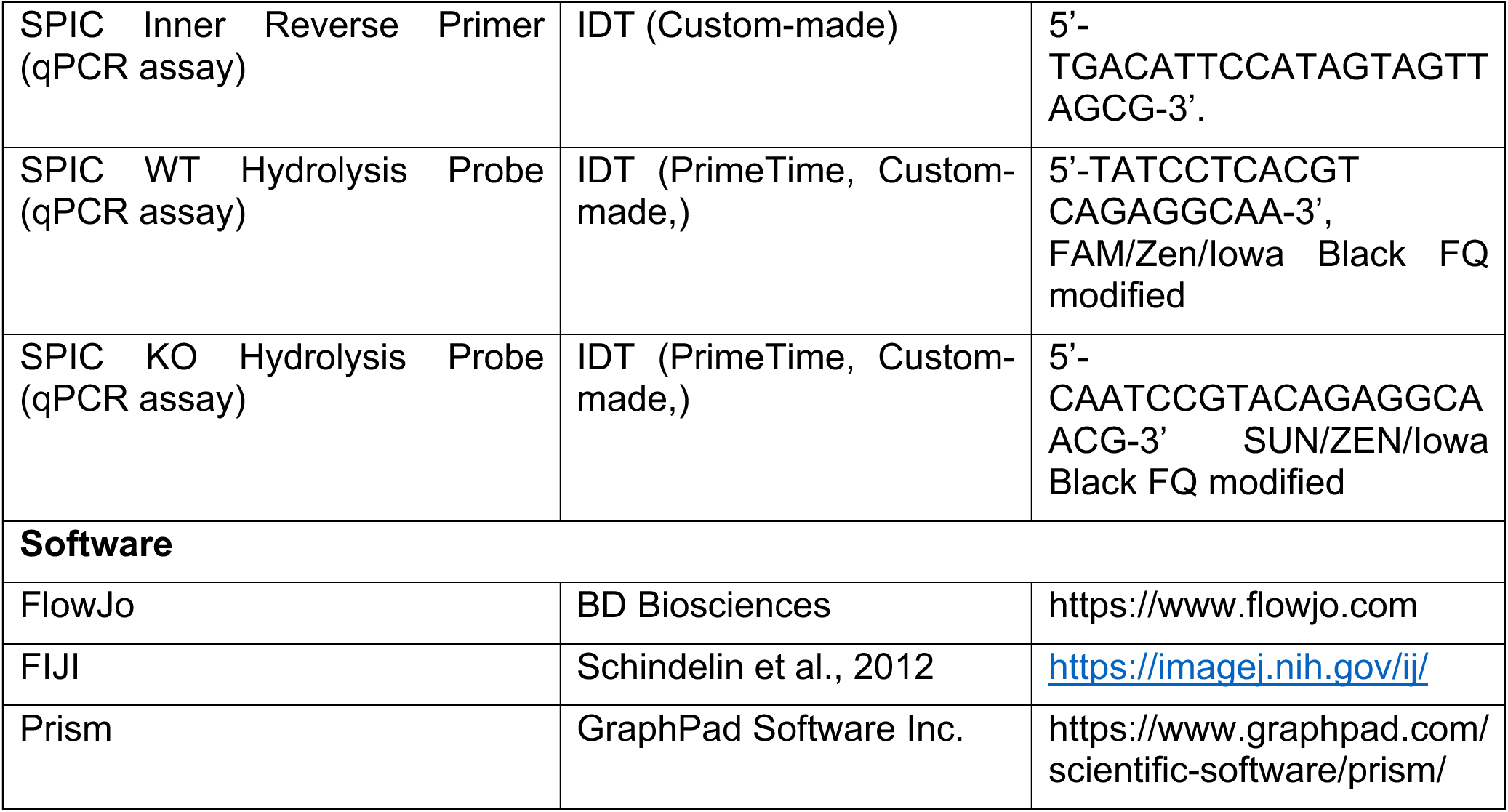

#### Immunofluorescence Microscopy

A full list of antibodies used for immunohistochemistry can be found the Key Resources table. Spleens were harvested, frozen in OCT compound (Fisher Scientific), and frozen sections 8 μm in thickness were placed on SuperFrost Plus cryosection slides (Fisher Scientific). Sections were fixed in ice-cold acetone at -20 °C for 10 minutes and then allowed to dry. A boundary was drawn around tissue sections using a pap pen (Fisher Scientific). Sections were washed with PBS and then blocked with staining buffer (PBS with 3% bovine serum albumin, ± 1% saponin, 5 % normal mouse serum) for 30 min at room temperature. After blocking, sections were stained with the primary anti-*Salmonella* antibody in staining buffer for 4 hours, or with primary antibodies against surface antigens for 1.5 hours, at room temperature. Sections were washed and then stained for 2 hours at room temperature with fluorescent conjugated secondary antibodies. Slides were washed in PBS and then mounted using ProLong Diamond (Life Technologies). Images were acquired on a Zeiss LSM 700 confocal microscope with the ZEN 2010 software (Zeiss) and processed using FIJI software.

#### Histology

Tissues were fixed in 10% neutral buffered formalin. After fixation, tissues were routinely processed for paraffin embedding and stained with hematoxylin and eosin. Images were collected using a ImageScope scanner and processed with Qupath software.

#### Cell preparation for 10X Genomics scRNA-sequencing

All samples subjected to scRNA-seq were prepared and FACS-enriched using the following procedures. *S*Tm infected spleens were harvested and digested in buffer containing HBSS + Ca^2+^ + Mg^2+^ + 50 μg/mL DNase (Roche) + 25 μg/mL Liberase TL (Sigma) at 37 °C for 25 min, mixing at 200 rpm. EDTA was added to 5 mM final concentration to stop digestion reaction and cells were washed with RPMI containing 10% FCS. Following red blood cell lysis, splenocytes were washed twice RPMI containing 10% FCS. Splenocytes were stained with an antibody mixture for surface markers (CD11b Alexa fluor 647, CD11c PECy7, Ly6C PerCP Cy5.5, Ly6G FITC, CD3 APC efluor 780, CD19 APC efluor 780, and NK1.1 APC efluor 780) for 25 minutes on ice, washed twice with RPMI containing 10% FCS, then resuspended in the same buffer with 1:2000 DAPI. Splenocytes were then FACS-enriched on a BD FACSAria cell sorter. A permissive gating strategy was utilized to simultaneously enrich CD11b^+^CD11c^+^Ly6C^+^ macrophages and capture other splenocytes for sequencing. Sorting gates were set tightly for size/scatter, singlet, and living cells but more loosely for CD11b^+^, Ly6G^-^, Ly6C^+^, and CD11c^+^ cells as shown in Figure S1. The viability of sorted cells was checked using Trypan blue staining and hemocytometer inspection under a light microscope. Samples had viability greater than 90%. Cells were resuspended to a concentration to 500-1200 cells/uL, partitioned, and captured for sequencing on a 10x Chromium Controller. Libraries were prepared by the Stanford Functional Genomics Facility (SFGF) using 10X Genomics 3’ kit and sequenced on the Illumina HiSeq4000 or NovaSeq platform to a depth of ∼40,000 - 50,000 reads/cell. Raw sequencing data were demultiplexed by SFGF to yield fastqs reads.

#### Sequencing alignment and data preprocessing

Paired-end reads were mapped to the *Mus musculus* genome reference GRCm38 using 10X Cell Ranger (version 7.1.0) with parameters “--expect-cells=10000 --chemistry=auto,” yielding UMI count matrices that were used directly for all downstream preprocessing and analyses. To eliminate low-quality cells, we kept cells with more than 500 detected genes and more than 1,000 UMI read counts, and removed stressed cells in which greater than 5% of total UMI counts mapped to the mitochondrial genome. Cells that passed these filters were merged across batches (n = 7 WT infected and uninfected libraries) for downstream time-course analysis, yielding 32,297 cells across 32,285 mouse genes. UMI counts were normalized for sequencing coverage such that each cell had a total count equal to the median library size across all cells, yielding counts per median (CPM). Normalized CPM were offset by a pseudocount of 1 and log2-transformed. Data preprocessing and transformation were performed using the Scanpy package (v1.8.2; Python 3.7).

#### Cell clustering and cell type annotation

Cell type identities for the cells profiled in the present study were transferred from the annotated reference dataset previously published by Pham et al. (2023)^42^. To minimize confounding from sex-linked transcriptional programs, we excluded chromosome X and chromosome Y genes (1,829 genes, identified from the GRCm38 GTF) and renormalized the gene count table prior to clustering. The self-assembling manifold (SAM) algorithm (v0.8.5) was then run on the expression matrix with a low-expression threshold of 0.02, a fixed random seed (seed = 123), and otherwise default parameters, producing gene weights, a nearest-neighbor graph, and a two-dimensional UMAP projection. Leiden community detection was performed at resolution = 0.75 to assign cluster membership. Reference cell type labels from Pham et al. (2023)^42^ were propagated to newly profiled cells through five iterations of nearest-neighbor majority voting on the SAM-inferred connectivity graph. Cells assigned to the mononuclear phagocyte populations (MoMac 1 and MoMac 2) were retained for monocyte/macrophage subclustering. After excluding a small set of pre-identified outlier cells, the MoMac subset comprised 9,011 cells, which were re-clustered with the same SAM and Leiden parameters to yield 14 leiden clusters used for subsequent marker gene and pathway analyses.

#### Differential gene expression analysis

Differential gene expression was computed between each leiden cluster and the union of all other clusters in the MoMac subset. To prevent sex-linked genes from artefactually appearing as cluster markers, chromosome X and chromosome Y genes (1,829 total) were excluded prior to differential testing, restricting the test space to 30,456 genes. A negative binomial likelihood-ratio test was then conducted on the raw UMI counts without the sex chromosome genes, which served as the primary statistical test for identifying differentially expressed genes. False discovery rate (FDR) for all tests was controlled using the Benjamini-Hochberg procedure. For the negative binomial test, genes were considered differentially expressed (DEGs) when they satisfied FDR < 0.05, log2 fold change > 0.5, and detection in at least 10% of cells in the query cluster.

#### Gene sets over-representation analysis

To identify functional pathways that were enriched more than expected by chance, we performed gene set over-representation analysis (GSOA) on the lists of cluster-specific DEGs. Only DEGs with FDR < 0.05, log2 fold change > 0.25, and detection in greater than 25% of cells in the query cluster were used as input. We measured the fraction of input DEGs overlapping each pathway in the MSigDB Hallmark gene sets (retrieved with the gseapy package, v1.0.4) and assessed enrichment significance with a hypergeometric test against a background equal to the total number of mouse genes annotated in GRCm38 (31,053 genes). FDR was again controlled with the Benjamini-Hochberg procedure, and gene sets with FDR < 0.1 were considered statistically significant.

#### Pathway activity score analysis

Per-cell pathway activity scores were computed using Scanpy’s sc.tl.score_genes function, which calculates, for each cell, the mean expression of an input gene set minus the mean expression of a size-matched reference set of genes binned by expression level. A positive score indicates higher expression of the pathway genes than expected from background. Eight gene sets were scored in the MoMac subset of this study: a red-pulp macrophage (SpiC-RPM) signature (Spic, Hmox1, Vcam1, Mertk, Hba-a1, Alas2, Fech, Slc40a1); an ACE^+^ macrophage signature (Ace, Eno3, S1pr5, Id3); a curated heme biosynthesis/degradation gene set comprising heme biosynthetic enzymes (Alas1, Alas2, Alad, Cpox, Ppox, Urod, Uros, Fech), heme catabolic enzymes (Hmox1, Hmox2, Blvra, Blvrb), and hemoglobin chains (Hba-a1, Hba-a2, Hbb-bs, Hbb-bt); and five MSigDB Hallmark gene sets retrieved from the MSigDB_Hallmark_2020 mouse library via gseapy (UV response up, TNFα signaling via NF-κB, interferon-α response, complement, and glycolysis). The MSigDB Hallmark “heme metabolism” set was deliberately replaced by the curated heme biosynthesis/degradation gene set above because the Hallmark version consists of a broad erythroid-program signature that is dominated by baseline-high maintenance genes whose expression decreases during infection-induced macrophage activation, in opposition to the canonical heme enzymes.

#### Quantification and Statistical Analysis

Sample sizes were chosen based on previous experience to obtain reproducible results. On rare occasions, an infected mouse within a cohort developed signs of meningitis or hemoperitoneum characterized by ruffling, ataxia, inability to attain water and food, and hunching. Such animals were humanely euthanized and excluded from analyses. Data were consistently reproduced in at least 2 independent experiments, with a minimum of 4 mice analyzed per group in each experiment, as indicated in the legends. All statistics were calculated in GraphPad Prism v8.0 software using two-tailed Mann-Whitney tests and are noted in figure legends. P values are expressed as follows: * p < 0.05, ** p < 0.01, *** p < 0.001, **** p < 0.0001.

## Data and Code Availability

All data needed to evaluate the conclusions in the paper are present in the paper and/or the Supplementary Materials. Dataset and analysis code can be found at TBD repository and Github.

