## Supplemental Materials for "SPIC-dependent erythrophagocytic macrophages drive granuloma formation and pathogen persistence during intracellular bacterial infection"

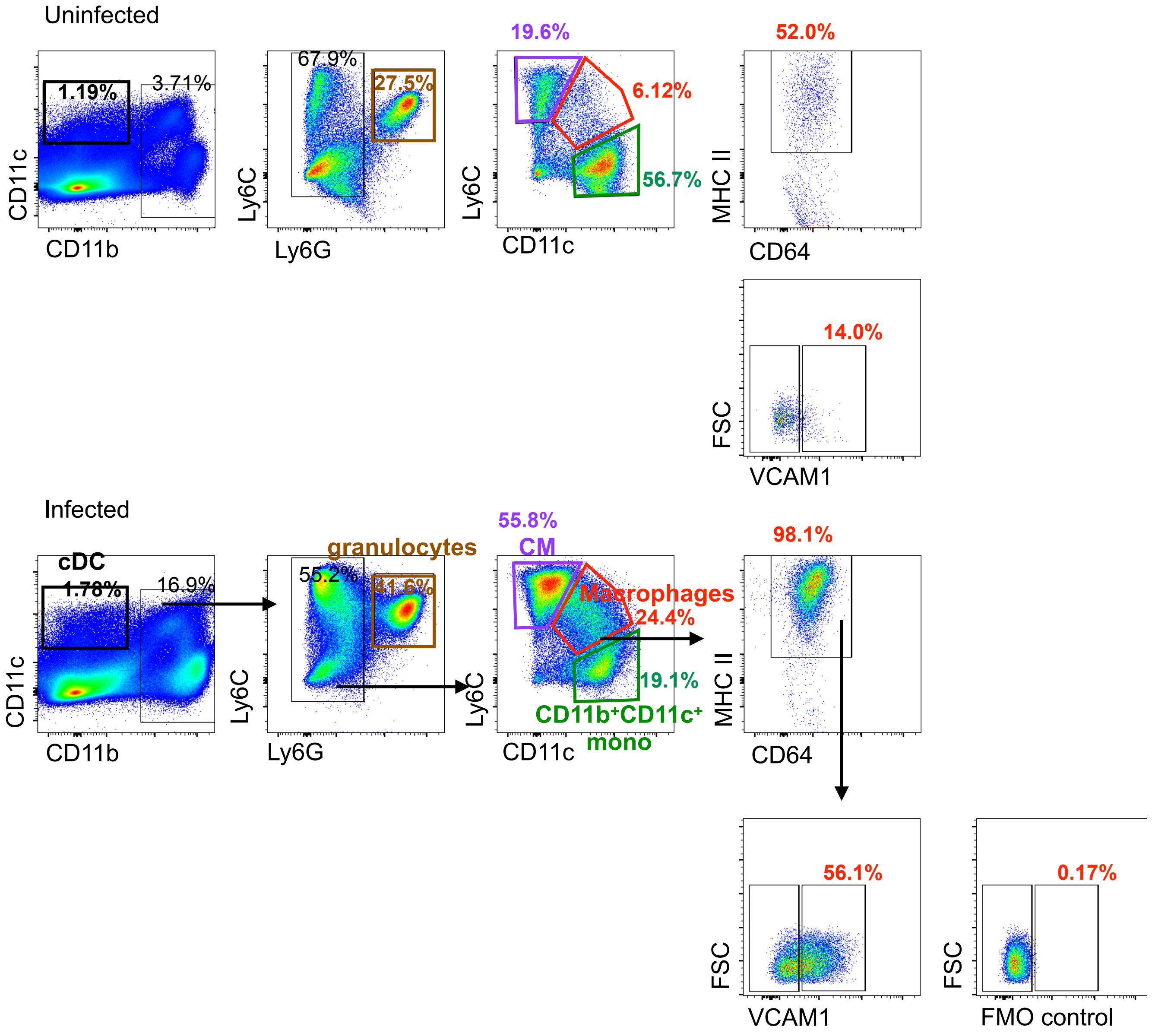


**Suppl. Figure 1**

**Figure S1. Related to Figure 2-6:** Gating scheme to identify and quantify CD11b$^{+}$CD11c$^{+}$Ly6C$^{+}$ granuloma-associated macrophages and other indicated myeloid cell populations. Erythrocyte-lysed splenocyte samples were stained and analyzed on flow cytometers. Samples were gated for size/scatter, singlet, and living population then further gated as displayed. For comparisons between SPIC WT and KO mice, the total gated macrophage populations are used for quantitating VCAM1, CD86, intracellular TER119, *S*Tm, CXCL9, and other phenotypic and functional marker differences.

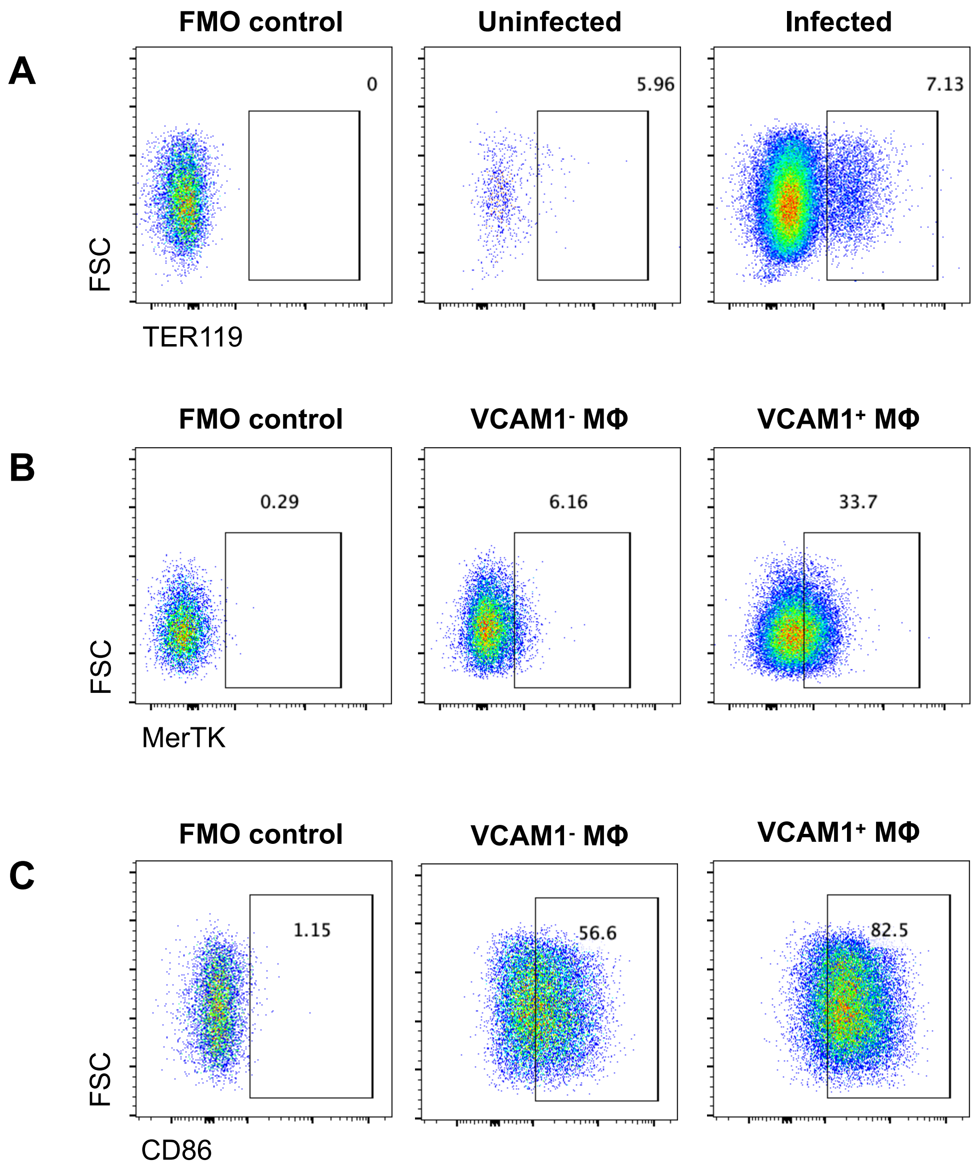


**Suppl. Figure 2**

**Figure S2. Related to Figure 2-5:** Gating scheme to identify and quantify functional properties of CD11b$^+$CD11c$^+$Ly6C$^+$ granuloma-associated macrophages and other indicated myeloid cell populations. Erythrocyte-lysed splenocyte samples were stained and analyzed on flow cytometers. Samples were gated for macrophages, classical monocytes (CM), classical dendritic cells (cDC), CD11b$^+$CD11c$^+$ monocytes, and in some analyses for VCAM1$^-$ and VCAM1$^+$ macrophages, as shown in Figure S1. (A) Gating for TER119$^+$ macrophages from uninfected and infected spleens. FMO: Fluorescence-Minus-One negative stain control. (B) Gating for MerTK$^+$ cells between VCAM1$^-$ and VCAM1$^+$ macrophages. FMO: Fluorescence-Minus-One negative stain control. (C) Gating for CD86$^+$ cells between VCAM1$^-$ and VCAM1$^+$ macrophages. FMO: Fluorescence-Minus-One negative stain control.

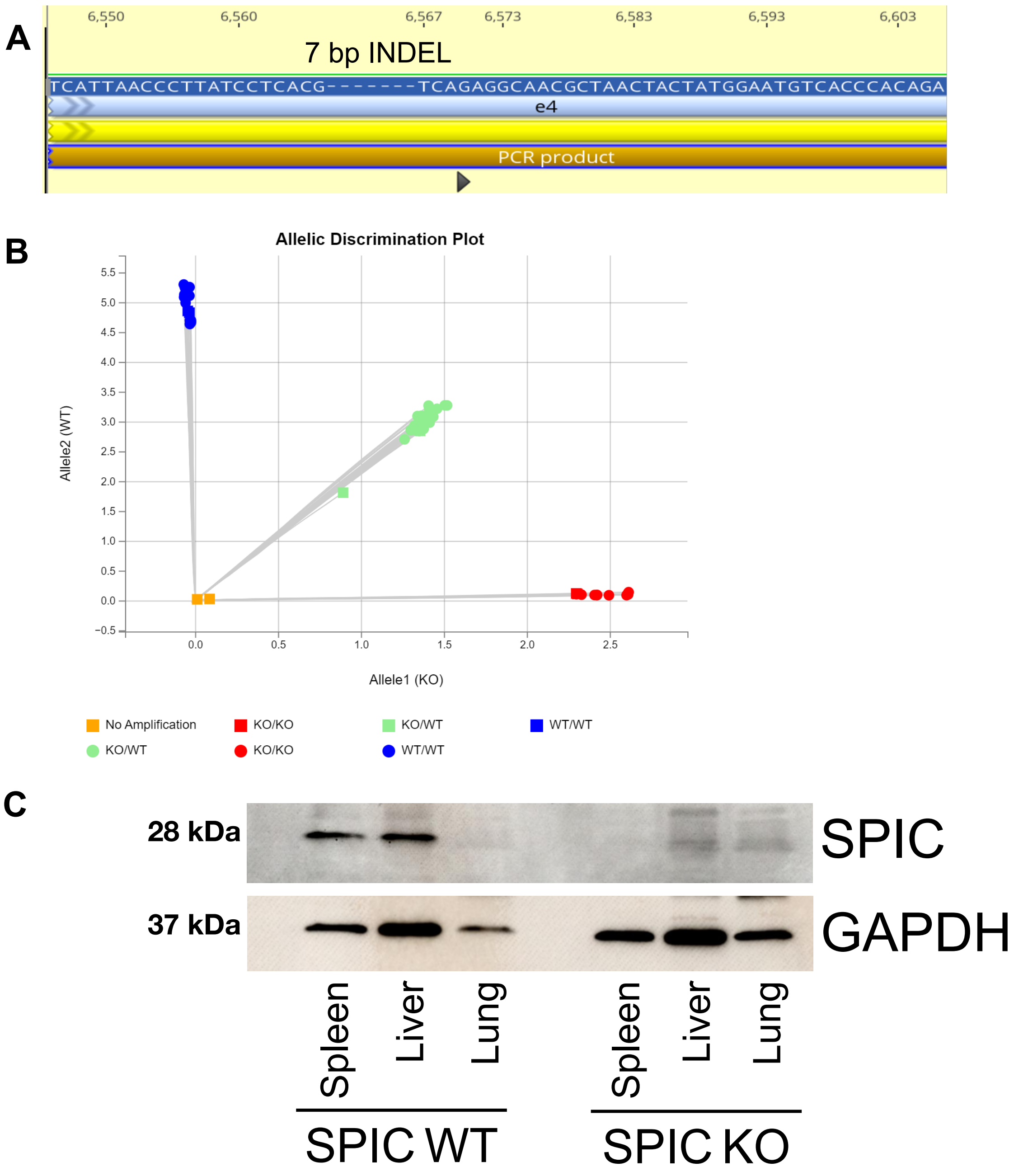


Suppl. Figure 3

**Figure S3. Related to Figure 4:** (A) Diagram showing the genomic position of the 7 nucleotide INDEL within Exon 4 of *Spic* of SPIC KO 129x1/SvJ mouse strain generated from CRISPR gene-editing. Black arrowhead indicates gRNA position. (B) Example of our *Spic* genotyping qPCR assay discriminating between the *Spic* wildtype and CRISPR-edited *Spic* KO allele. (C) Western blot analysis for SPIC expression with SPIC-WT and KO mice.

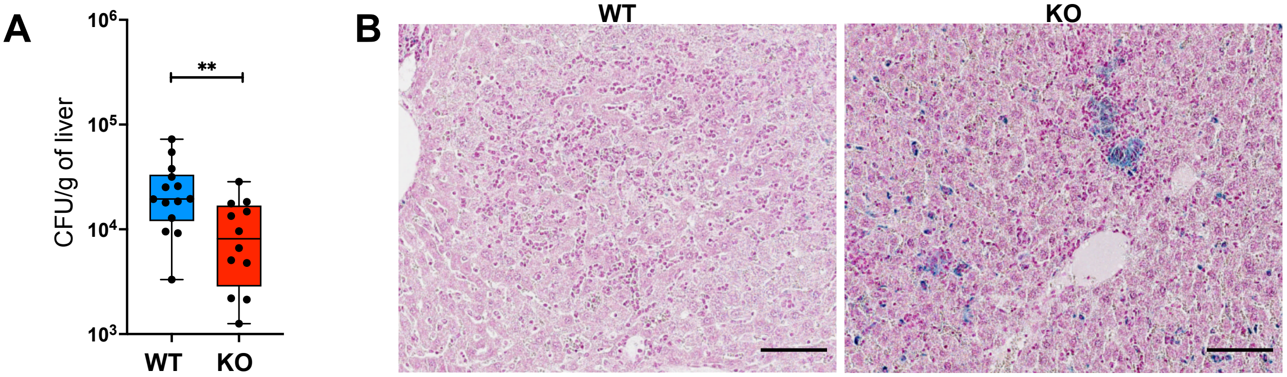


Suppl. Figure 4

**Figure S4. Related to Figure 4:** (A) Bacterial levels in infected SPIC WT and KO livers quantitated by CFU plating assay. Dots: individual mice. Data from 3 independent experiments 4-5 WT and KO mice per experiment. (B) Perl's Prussian Blue stain for ferric iron on formalin-fixed livers from infected SPIC WT and KO mice. Scale bar: 100 $\mu$m. Significance calculated using a two-tailed Mann-Whitney test.

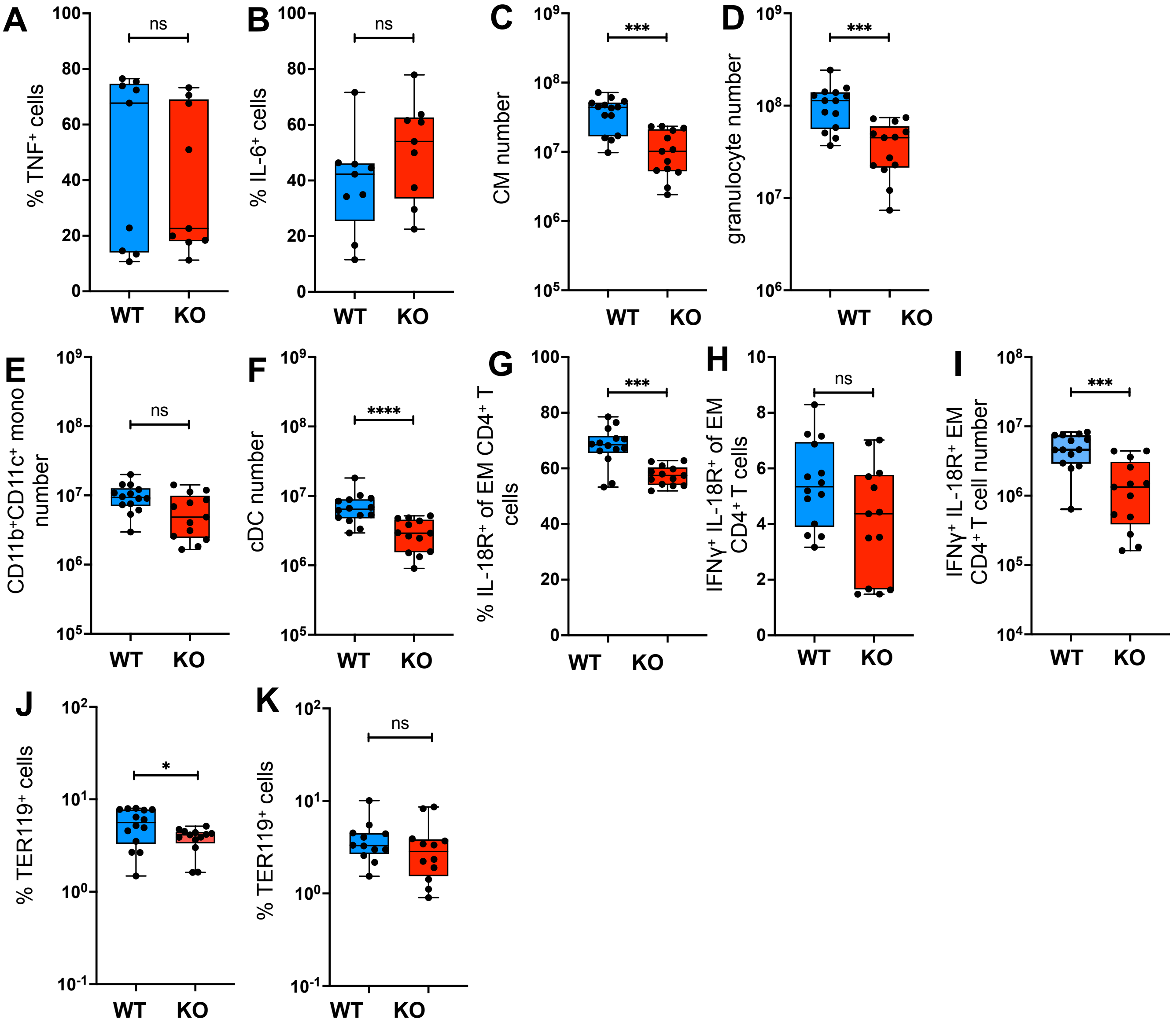


Suppl. Figure 5

**Figure S5. Related to Figure 5:** (A-I) Infected mice analyzed at 4 weeks p.i. (A and B) Splenocytes from infected spleens were stimulated with heat-killed *S*Tm ex-vivo and intracellular cytokine levels were quantitated using flow cytometry analysis (Methods). Samples gated for macrophages and other myeloid cells as shown in Figure S1. Frequencies of intracellular $TNF^+$ and $IL\text{-}6^+$ levels of macrophages shown. (C) Numbers of classical monocytes (CM) in SPIC WT and KO spleens. (D) Numbers of granulocytes in SPIC WT and KO spleens. (E) Numbers of $CD11b^+CD11c^+$ monocytes in SPIC WT and KO spleens. (F) Numbers of classical dendritic cells (cDC) in SPIC WT and KO spleens. (G) Frequencies of $IL\text{-}18R^+$ among effector memory $CD4^+$ T cells in SPIC WT and KO mice. Cells are for $CD3^+CD4^+CD62L^-CD44^+$ before plotting for IL-18R. (H) Frequencies of $IFN\gamma^+$ among $IL\text{-}18R^+$ effector memory $CD4^+$ T cells (Th1 cells) in SPIC WT and KO mice. Cells are for $CD3^+CD4^+CD62L^-CD44^+$ before plotting for IL-18R and IFNγ. (I) Numbers of $IFN\gamma^+IL\text{-}18R^+$ effector memory $CD4^+$ T cells (Th1 cells) in SPIC WT and KO mice. Cells are for $CD3^+CD4^+CD62L^-CD44^+$ before plotting for IL-18R and IFNγ. (J) Frequencies of $TER119^+$ macrophages in infected WT and KO spleens at 4 weeks p.i. (K) Frequencies of $TER119^+$ macrophages in infected WT and KO spleens at 2 weeks p.i. Significance calculated using a two-tailed Mann-Whitney test. A-I, data from 3 independent experiments 4-5 WT and KO mice per experiment.

SPIC WT 1 SPIC WT 2 SPIC WT 3

SPIC KO 1 SPIC KO 2 SPIC KO 3

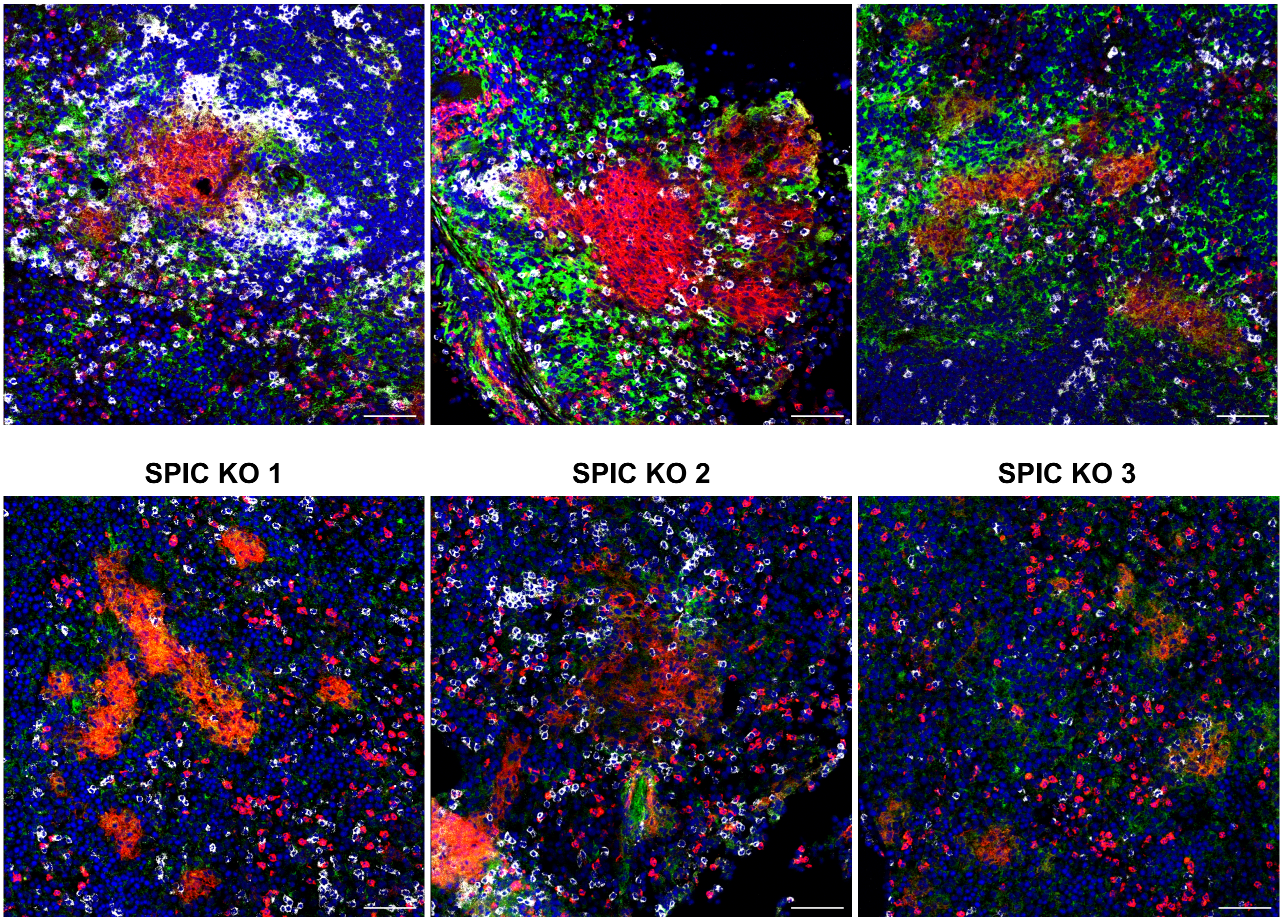

VCAM1 CD11b. CD4 DAPI

Suppl. Figure 6

**Figure S6. Related to Figure 6:** Additional confocal images showing impaired VCAM1$^+$ macrophage-T cell interaction zone at the granuloma periphery and loss of concentration of CD4 T cells in the lymphocytic cuffs at 4 weeks p.i.
